# Enteric glial cells favour accumulation of anti-inflammatory macrophages during the resolution of muscularis inflammation

**DOI:** 10.1101/2021.06.10.447700

**Authors:** Michelle Stakenborg, Saeed Abdurahiman, Veronica De Simone, Gera Goverse, Nathalie Stakenborg, Lies van Baarle, Qin Wu, Dimitri Pirottin, Jung-Seok Kim, Louise Chappell-Maor, Isabel Pintelon, Sofie Thys, Louis Boon, Marlene Hao, Jo A. Van Ginderachter, Guy E. Boeckxstaens, Jean-Pierre Timmermans, Steffen Jung, Thomas Marichal, Sales Ibiza, Gianluca Matteoli

**Author notes:** these authors contributed equally to this work (Michelle Stakenborg, Saeed Abdurahiman). senior authors: Sales Ibiza and Gianluca Matteoli. Corresponding author: Prof. Sales Ibiza, PhD Laboratory of Cell Biology & Histology Department of Veterinary Sciences University of Antwerp, Campus Drie Eiken, Universiteitsplein 1, B-2610 Wilrijk, Belgium Prof. Gianluca Matteoli, DVM, PhD Laboratory of Mucosal Immunology Department of Chronic Diseases, Metabolism and Ageing (CHROMETA) KU Leuven, Herestraat 49, O&N1 box 701| BE-3000 Leuven | Belgium.

## Abstract

**Objective:** Monocyte-derived macrophages (Mφs) are crucial regulators during muscularis inflammation. However, it is unclear which microenvironmental factors are responsible for monocyte recruitment and neurotrophic Mφ differentiation in this paradigm. Here, we investigate Mφ heterogeneity at different stages of muscularis inflammation and determine how environmental cues can attract and activate tissue protective Mφs.

**Design:** Single cell RNA sequencing was performed on immune cells from the muscularis of wild-type and CCR2^-/-^ mice at different timepoints after muscularis inflammation. CX3CR1^GFP/+^ and CX3CR1^CreERT2^ R26^YFP^ mice were analyzed by flow cytometry and immunofluorescence. The transcriptome of enteric glial cells (EGCs) was investigated using PLP^CreERT2^ Rpl22^HA^ mice. In addition, we assessed the effect of supernatant from neurosphere-derived EGCs on monocyte differentiation based on the expression of pro- and anti-inflammatory factors.

**Results:** Muscularis inflammation induced marked alterations in mononuclear phagocyte populations associated with a rapid infiltration of Ly6c^+^ monocytes that locally acquired unique transcriptional states. Trajectory inference analysis revealed two main pro-resolving Mφ subpopulations during the resolution of muscularis inflammation, *i.e*. Cd206^+^ MhcII^hi^ and Timp2^+^ MhcII^lo^ Mφs, which were both derived from CCR2^+^ monocytes. Interestingly, we found that EGCs were able to sense damage to the muscularis to stimulate monocyte recruitment and differentiation towards pro-resolving Mφs via CCL2 and CSF1, respectively.

**Conclusion:** Our study provides a comprehensive insight into pro-resolving Mφ differentiation and their regulators during muscularis inflammation. We deepened our understanding in the interaction between EGCs and Mφs, thereby highlighting pro-resolving Mφ differentiation as a potential novel therapeutic strategy for the treatment of intestinal inflammation.

## Introduction

In recent years, significant advances have been made in our understanding of the phenotype of intestinal macrophages (Mφs), which perform niche-defined functions to maintain immune homeostasis and regulate motility and secretion via the enteric nervous system (ENS)^1–3^. Consequently, specific niches are populated by different Mφ subsets with a unique transcriptome to meet the diverse functional demands of the tissue microenvironment. For instance, lamina propria Mφs reside in close proximity to the epithelium and at the crypt base, where they are involved in the maintenance of gut homeostasis by contributing to epithelial barrier integrity and by phagocytosing luminal antigens^2–5^. In the deeper layers of the gut wall at the level of the muscularis externa, Mφs exert neurotrophic functions to promote enteric neuron survival and to support neuronal function^1, 6, 7^. However, during muscularis inflammation, activation of resident Mφs results in impairment of neuromuscular function due to the release of nitric oxide, prostaglandins and cytokines, simultaneously triggering the influx of inflammatory cells^8–10^. Although we have shown in a previous study that the influx of monocytes is essential for the resolution of muscularis inflammation due to their differentiation into pro-resolving monocyte-derived Mφs^11^, it remains to be determined which cell types are responsible for monocyte recruitment and which factors promote their differentiation towards anti-inflammatory Mφs during muscularis inflammation. The extensive crosstalk between muscularis Mφs and the ENS at homeostasis suggests a potential involvement during inflammation^6^. The ENS is organized in a network of ganglia containing neurons and enteric glial cells (EGCs). Historically, EGCs were mainly considered as supporting cells of enteric neurons. However, recent evidence suggests that EGCs have a much broader function in gastro-intestinal (GI) physiology, contributing to motility and preserving epithelial barrier integrity by the interaction with innate lymphoid cells (ILCs), interstitial cells of Cajal, endothelial and epithelial cells^12–17^. Even more intriguing is the putative role of EGCs in immune regulation. In this regard, the interaction between astrocytes and microglia has been extensively studied in the brain during development, homeostasis and disease^18^. Yet, limited studies have investigated the interaction between Mφs and EGCs in the intestine^19^.

Here, we investigate Mφ heterogeneity and their role in tissue repair during muscularis inflammation. Using time-series single-cell transcriptomics, we observed that muscularis inflammation induced a prominent myeloid cell diversification, resulting in 7 myeloid subpopulations. Trajectory inference analysis indicated that incoming Ly6c^hi^ monocytes acquired diverse gene expression signatures in the injured muscularis, notably resulting in two main pro-resolving Mφ subpopulations characterized by Cd206^+^ MhcII^hi^ and Timp2^+^ MhcII^lo^ gene signatures. In addition, EGCs were identified as the main producers of the monocyte chemo-attractant, CCL2, during the early phase of inflammation. Furthermore, these cells produced a multitude of secreted factors that can potentially stimulate the differentiation of monocytes into pro-resolving Mφs. Specifically, we defined EGC-derived CSF1 as a critical factor for the generation of anti-inflammatory CD206^+^ Mφs *ex vivo*. Overall, our findings imply that during intestinal inflammation, future therapies should be aimed at enhancing the pro-resolving and neurotrophic properties of Mφs to reduce possible damage to the ENS and promote GI recovery.

## Material & methods

### Mouse Model

Twelve week old female wild-type (WT; C57BL/6J), CCR2 knockout (CCR2^-/-^)^20^, Cx3cr1^GFP/+, 21^, Cx3cr1^CreERT2, 22^ Rosa26-LSL-YFP^23^, PLP^CreERT2, 24^ Rpl22^HA, 25^ and their littermate controls were used. All lines were bred and maintained at the KU Leuven animal facility on a 12:12-h light-dark cycle and had *ad libitum* access to tap water and commercially available chow (ssniff® R/M-H, ssniff Spezialdiäten GmbH). All experimental procedures were approved by the Animal Care and Animal Experiments Ethical Committee of KU Leuven.

### Experimental model of small intestinal inflammation

An established model of intestinal manipulation was used to induce small intestinal muscularis inflammation as previously reported^26^. In brief, animals were anesthetized by intraperitoneal injection of ketamine (Ketalar 100 mg/kg; Pfizer) and xylazine (Rompun 10 mg/kg; Bayer). After a midline laparotomy, the cecum and the small intestine were carefully externalized and mounted onto a plexiglas platform. Next, the small intestine was manipulated three times back and forth using a purpose-designed device that enables the application of a constant pressure to the intestine by a cotton swab applicator attached to its end.

### Statistical analysis

Results are shown as mean ± standard error of the mean (SEM). Significance between two mean groups was determined by an unpaired two-tailed Student’s t-test or a non-parametric Mann-Whitney test, while one-way analysis of variance (one-way ANOVA) followed by Dunnett’s Multiple comparison test was performed to compare the mean of multiple groups. GraphPad Prism V.9.1.0 software (GraphPad Inc) was used to generate graphs and to perform statistical analysis.

## Results

### Time-dependent recruitment of immune cell in the inflamed muscularis

Surgery-induced damage to the muscularis leads to a transient impairment of GI motility associated with extensive recruitment of immune cells to the ENS. To characterize the immune cell infiltrate and their respective activation states during muscularis inflammation, droplet-based scRNA-seq was performed on sorted CD45^+^ immune cells from the muscularis of naïve mice, and during the acute (24h) and resolving phase of muscularis inflammation (72h) (10X Genomics Platform; Fig. 1A). Unsupervised clustering of 4,102 cells and reference-based identification using Immgen (Fig. 1B-C **and** Supplementary Fig. 1A-B) revealed 12 independent immune cell populations including monocytes (*Ccr2, Ly6c2, Chil3*), 3 clusters of Mφs (1: *Cd63, Cd68, Trem2*; 2: *Itgam, Arg1, Lyz2*; 3: *Cx3cr1, Csf1r, Mrc1*), dendritic cells (DCs; *Itgax, Cd209a, Ccl17*), 2 clusters of neutrophils (1: *Cxcr2, Mmp9, S100a9*; 2:, *S100a8, Hcar2, Msrb1*), eosinophils (*Siglec-f, Cxcr4, Pim1*), ILCs (*Thy1, Rora, Il7r*), T cells (*Cd3e, Cd7, Trdc*) and 2 clusters of B cells (1: *Cd79a, Cd20, CD79b*; 2: *Eaf2, Mef2b, CD19*)^27^.

**Figure 1:**
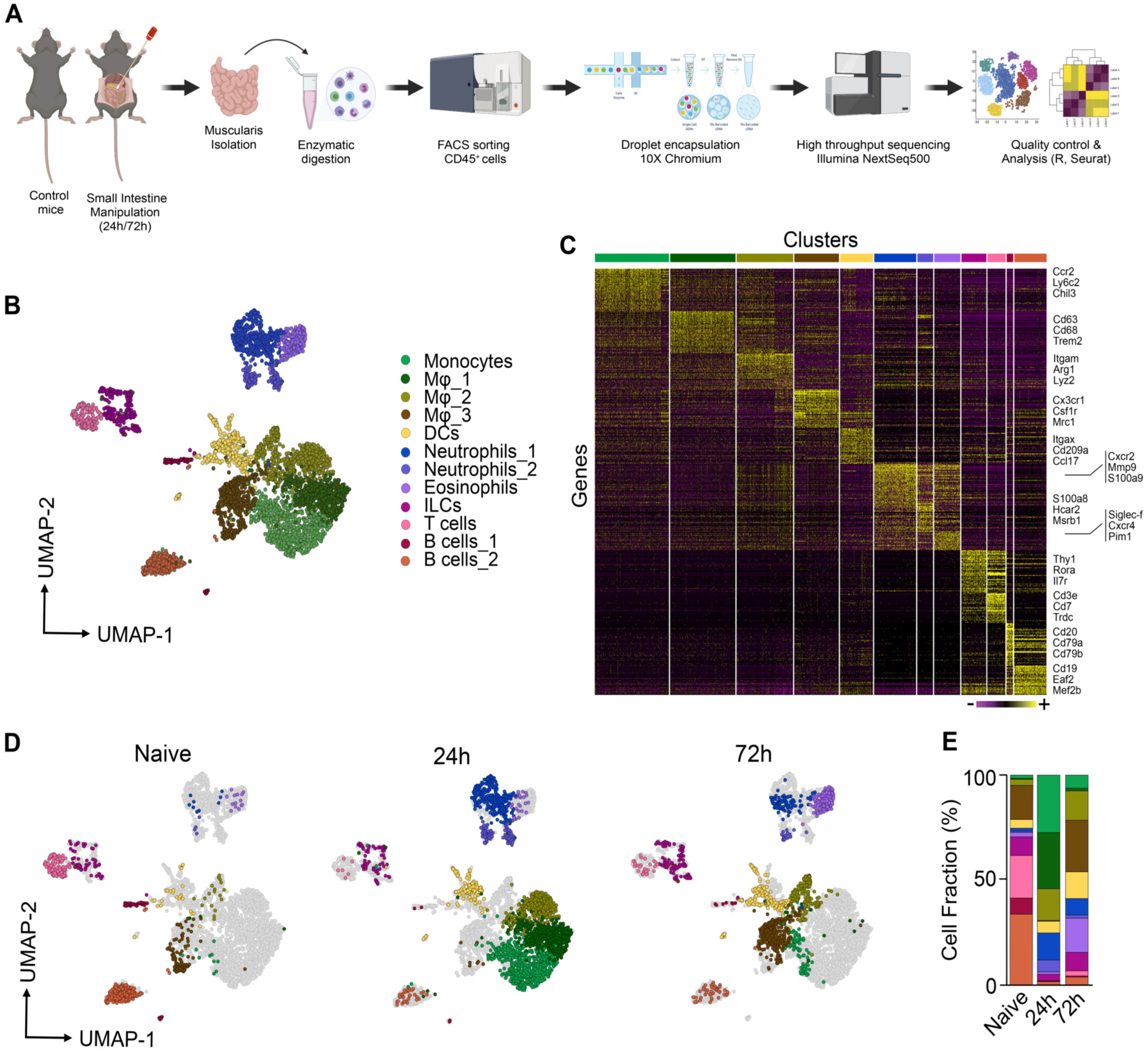
Identification of CD45^+^ immune cell populations by unsupervised scRNAseq clustering in the healthy and inflamed muscularis. **A.** Experimental pipeline of scRNAseq experiment. **B.** UMAP of sorted CD45^+^ immune cells from the healthy muscularis, 24h and 72h post-injury from WT mice. Each sample was pooled from 3-4 mice. **C.** Heatmap of the 50 most differentially expressed genes in each cluster. **D.** UMAPs of time-dependent infiltration of immune cells upon muscularis inflammation. Data of Figure 1B is split out based on different time points. **E.** Cell fraction of each cluster relative to the total number of CD45^+^ immune cells at different time points after muscularis inflammation.

To examine the role of different immune cell populations, we determined their gene expression signatures at different time points during muscularis inflammation. At homeostasis, the muscularis mainly consisted of a population of Cx3cr1^+^ Mφs (Mφ_3) together with T cells, B cells and ILCs (Fig. 1D-E). In the acute and resolving phase of muscularis inflammation, and consistent with previous observations^11, 28^, a shift in the leukocyte populations with respect to the homeostatic condition was observed (Fig. 1D-E). In the acute phase (24h), there was a massive infiltration of monocytes in addition to Mφs (Mφ_1 and Mφ_2) and neutrophils (Neu_1 and Neu_2), pointing towards a more pro-inflammatory micro-environment. During the resolution of muscularis inflammation (72h), the muscularis was mainly populated by Cx3cr1^+^ Mφs (Mφ_3), DCs and eosinophils, indicating a return to homeostasis. Altogether, these results show that during muscularis inflammation a shift in the immune landscape favors the resolution of inflammation and a rapid return to homeostasis.

### Identification of two distinct myeloid subpopulations during the resolution of muscularis inflammation

Monocyte-derived Mφs are essential for the recovery of GI motility during the resolution of muscularis inflammation^11, 28^. However, their heterogeneity and differentiation trajectory towards tissue protective Mφs have not yet been characterized during muscularis inflammation. To this end, subsets originally identified as monocyte/Mφ subpopulations in our scRNAseq dataset (Fig. 1) were extracted and re-clustered to better define which type of myeloid cells might aid in the resolution of muscularis inflammation, thereby identifying 7 distinct subsets (Fig. 2A-D **and** Supplementary Fig. 2A). At homeostasis, the muscularis micro-environment mainly consisted of Cx3cr1^+^ Mφs with high expression of typical resident Mφ markers such as *Cd81*, *Cd72* and *H2-Eb1* (Fig. 2E). During acute muscularis inflammation, the most predominant subpopulation was the cluster of classical Ly6c^+^ monocytes, which was enriched for the expression of *Plac8*, *Hp* and *Chil3* (Fig. 2B-E). Additionally, a subpopulation with a gene expression signature suggestive of an intermediate monocyte-to-Mφ differentiation state was observed (Ccr2^+^ int Mφs), which was underscored by their moderate *MhcII* expression as compared to Ly6c^+^ monocytes and homeostatic Cx3cr1^+^ Mφs (Fig. 2B). Besides Ly6c^+^ monocytes and Ccr2^+^ int Mφs, we observed a subcluster of Arg1^+^ Mo/Mφ with high expression of *Ccl9* and *Srgn* alongside Fabp5^+^ Mo/Mφ with characteristic expression of *Lgals1*, *Ccl7* and *Flt1* (Fig. 2E). Of note, 72h post-injury, during the resolution of muscularis inflammation, Ly6c^+^ monocytes and Ccr2^+^ int Mφs were present at a low percentage but two novel Mφ subclusters were identified: Cd206^+^ Mφs and Timp2^+^ Mφs (Fig. 2C-D **and** Supplementary Fig. 2B). The most abundant subcluster during the resolution of muscularis inflammation, Cd206^+^ Mφs, displayed high *MhcII* expression similar to homeostatic Cx3cr1^+^ Mφs, in addition to high expression of anti-inflammatory genes such as *Selenop*, *Mrc1*, *Igf1*, *Trem2* and *Stab1* (Fig. 2B **and** Supplementary Fig. 2B-C). In contrast, Timp2^+^ Mφs had lower *MhcII* expression compared to homeostatic Cx3cr1^+^ Mφs, but similarly expressed high levels of tissue reparative markers such as *Ltc4s* and *Adgre5* (Fig. 2B, 2E **and** Supplementary Fig. 2B-C). To further investigate differences in the transcriptional regulation of Cd206^+^ and Timp2^+^ Mφs, single-cell regulatory network inference and clustering (SCENIC) was employed to assess specific regulon activities in different myeloid subclusters (Fig. 2F **and** Supplementary Fig. 2F). Cd206^+^ Mφs displayed high regulon activity of *FosB, Runx1* and *Irf8* similar to Cx3cr1^+^ Mφs, while Timp2^+^ Mφs had a distinctly altered transcription factor signature with high *Cebpb* and *Gata6* regulon activity.

**Figure 2:**
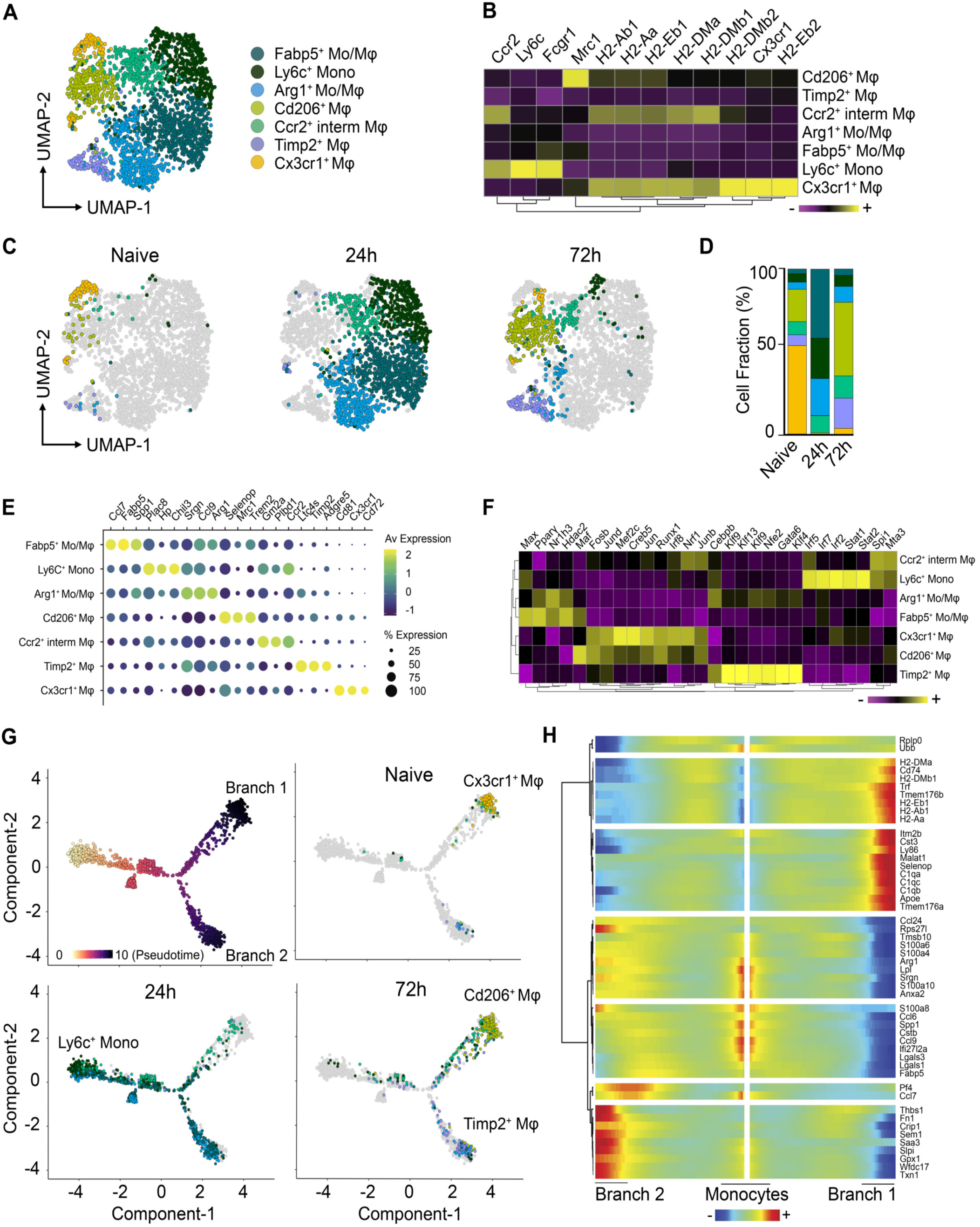
Two main anti-inflammatory Mφ subpopulations with a unique transcriptional state are present during the resolution of muscularis inflammation. **A.** UMAP of reclustered monocyte/Mφ subpopulations from Fig. 1A from the healthy muscularis, 24h and 72h post-injury from WT mice. **B.** Heatmap of typical monocyte and Mφ markers including MHCII genes. **C.** UMAPs of time-dependent infiltration of monocytes/Mφs upon muscularis inflammation. **D.** Cell fraction of each subcluster relative to the total number of monocytes/Mφs cells at different time points after muscularis inflammation. **E.** Dotplot showing expression of selected differentially expressed genes in each subcluster. **F.** Heatmap of regulon activity per cluster according to SCENIC analysis. **G.** Pseudotime analysis of monocytes/Mφs at different time points after muscularis inflammation. **H.** Heatmap of gene expression showing the top 50 genes of different branches of the pseudotime trajectory tree.

To evaluate whether these subclusters can also be detected by flow cytometry, a comparative study was performed during muscularis inflammation. In the healthy muscularis, only 1 subset of MHCII^hi^ Mφs was observed similar to homeostatic Cx3cr1^+^ Mφs (Supplementary Fig. 2D-E). During acute muscularis inflammation, the muscularis microenvironment was mainly dominated by infiltrating Ly6C^hi^ monocytes, which differentiated into Ly6C^+^ MHCII^+^ immature Mφs. However, at single cell level, we observed 4 different Mφ subclusters, indicating that Mφ heterogeneity was higher than originally considered. Interestingly, 72h post-injury, flow cytometric analysis also revealed only two main Mφ subclusters (MHCII^hi^ and MHCII^lo^ Mφs), similar to our observations at single cell level (Supplementary Fig. 2D-E).

Identifying relevant gene expression changes in myeloid subclusters during muscularis inflammation could shed light on the molecular mechanism regulating Mφ differentiation. In order to define these temporal transcriptional alterations leading to differentiation into Cd206^+^ and Timp2^+^ Mφs, Monocle-2 was used to superimpose subclusters on a trajectory placing Ly6c^+^ monocytes at the beginning of the pseudotime (Fig. 2G-H)^29^. We identified a major trajectory bifurcation leading to a branch of cells with high expression of genes found in homeostatic Mφs such as *Cd74*, *Tmem176a/b*, complement genes (*C1qa, C1qb, C1qc*) and *MhcII* genes (*H2-Eb1, H2-Ab1, H2-Aa*; Branch 1), while the other branch had less complement and *MhcII* expression with increased expression of *Fn1, Ltc4s* and *Saa3* (Branch 2) (Fig. 2H **and** Supplementary Fig. 2G-H). Interestingly, the Cd206^+^ and Timp2^+^ Mφ subsets were at two opposite ends of the trajectory suggesting alternative differentiation routes during the resolution of muscularis inflammation (Fig. 2G). Of note, homeostatic Cx3cr1^+^ Mφs were located in close proximity to Cd206^+^ Mφs in the pseudotime, underscoring the similarity between both Mφ subsets. Arg1^+^ and Fabp5^+^ Mo/Mφ were mainly present in Branch 2, while Ccr2^+^ int Mφs were exclusively present in Branch 1, indicating that these subpopulations might act as intermediates before attaining their terminal differentiation state. These findings support the hypothesis that there are two major monocyte-to-Mφ differentiation trajectories in the inflamed muscularis giving rise to Cd206^+^ and Timp2^+^ Mφs during the resolution of inflammation.

### Pro-resolving Mφs originate from CCR2^+^ monocytes during muscularis inflammation

In a previous study, we have shown that the influx of CCR2^+^ monocytes is essential for resolution of muscularis inflammation, as blocking monocyte infiltration resulted in a delayed recovery of GI motility and damage to the ENS^11^.To investigate whether CCR2^+^ monocytes are the source of pro-resolving Mφs during muscularis inflammation, scRNAseq was performed on sorted CD45^+^ immune cells from the muscularis of WT and CCR2^-/-^ mice 24h and 72h post-injury (Fig. 3A-C **and** Supplementary Fig. 3A-D). After extraction and re-clustering of the mono/Mφ subsets (2,406 cells), the gene expression signatures from the myeloid subsets in Fig. 2 were crossmatched by singleR with this novel dataset to specifically annotate the subpopulations (Supplementary Fig. 3C), yielding only one additional cluster, *i.e. Fn1*^+^ mono/Mφs. In CCR2^-/-^ mice, there was a large alteration in the myeloid compartment after muscularis inflammation. As expected upon acute muscularis inflammation in CCR2^-/-^ mice, there was an almost complete loss of Ly6c^+^ monocytes, and Arg1^+^ and Fabp5*^+^* Mo/Mφs. In addition, we could only detect a small subpopulation of Cx3cr1*^+^* Mφs representing the long-lived resident Mφs. During the resolution of muscularis inflammation, the Cd206^+^ and Timp2^+^ Mφ subsets were also completely absent in CCR2^-/-^ mice, confirming that these subpopulations are derived from CCR2^+^ monocytes and that they possibly represent the pro-resolving Mφs essential for recovery of GI motility after tissue damage.

**Figure 3:**
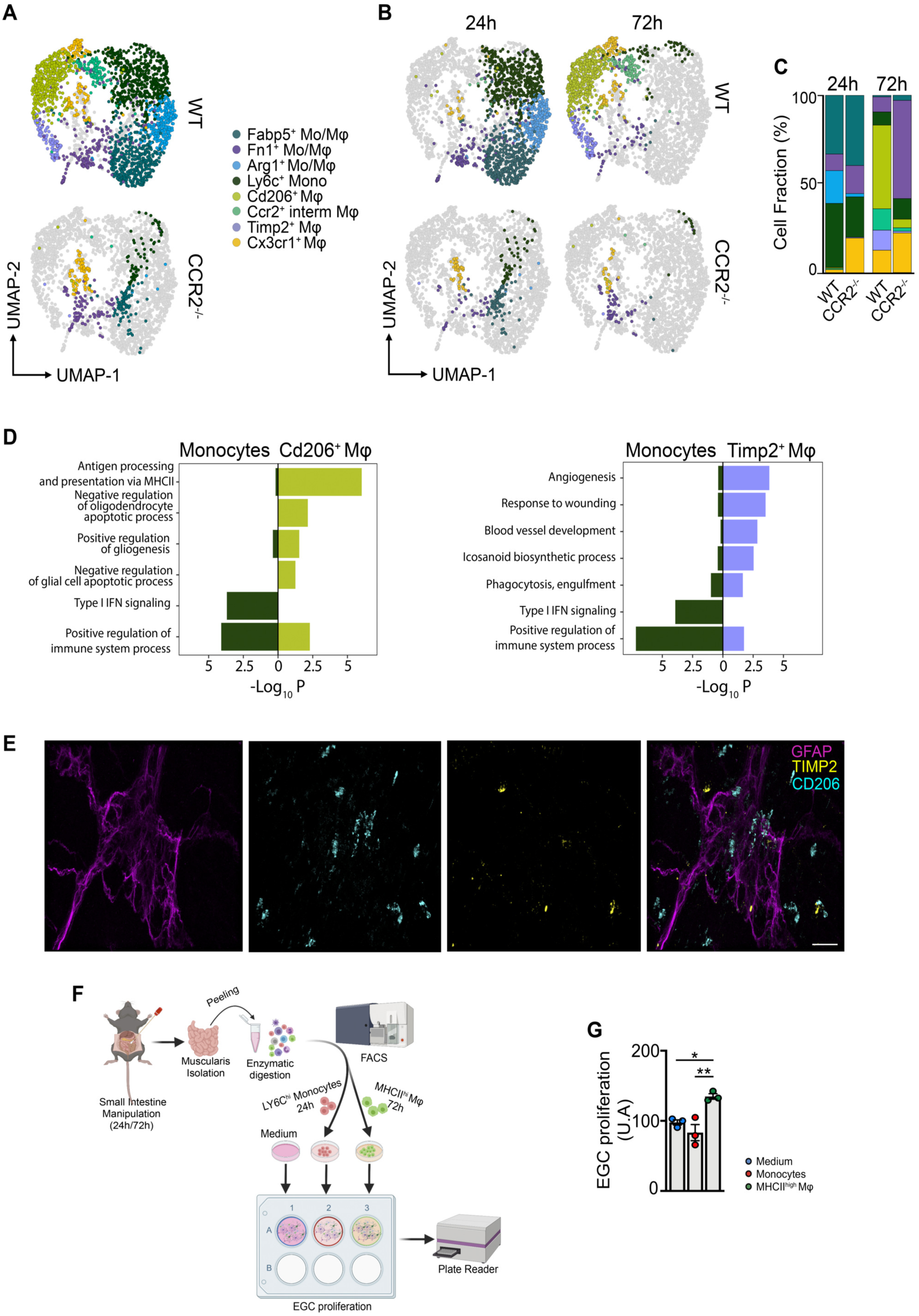
Two distinct Mφ subpopulations during the resolution of muscularis inflammation are derived from CCR2^+^ monocytes. **A.** UMAP of monocyte/Mφ subclusters from the muscularis of WT and CCR2^-/-^ mice at 24h and 72h after induction of muscularis inflammation. Each sample was pooled from 3-4 mice. **B.** UMAPs of myeloid cells at different time points post-injury. **C.** Cell fraction of each subcluster relative to the total number of myeloid cells at different time points after muscularis inflammation. **D.** GO analysis of monocytes versus Cd206^+^ Mφs (left) or Timp2^+^ Mφs (right) showing negative Log_10_(p-value). **E.** Immunofluorescent images of muscularis whole-mount preparations 3 days after the induction of muscularis inflammation stained for GFAP (purple), TIMP2 (yellow) and CD206 (light blue). Scale bar 15 µm. **F.** Experimental outline of *in vitro* EGC proliferation by stimulation with supernatant of monocytes/Mφs from different time points post-injury. Every data point is an independent sorting and culture experiment. **G.** Fold induction of EGCs stimulated with the supernatant of LY6C^hi^ monocytes from 24h post-injury or MHCII^hi^ Mφs from 72h post-injury relative to control medium. One-way ANOVA; test *p<0.05; **p<0.01; ns=not significant.

To further investigate the phenotype of Cd206^+^ and Timp2^+^ Mφs during muscularis inflammation, gene ontology (GO) analysis was performed and revealed that both Mφ subsets seemed pro-resolving in nature (Fig. 3D). For instance, Timp2^+^ Mφs were implicated in pathways associated with tissue repair and regulation of the inflammatory response, such as angiogenesis, response to wounding, blood vessel development, icosanoid biosynthetic processes, and phagocytosis-engulfment. Conversely, Cd206^+^ Mφs were enriched for pathways associated with antigen processing and presentation via MHC class II, negative regulation of oligodendrocyte and glial cell apoptotic processes and positive regulation of gliogenesis. In contrast, monocytes were clearly enriched for pro-inflammatory pathways, such as type I interferon signaling and positive regulation of immune system processes. Interestingly, CD206^+^ Mφs were located in close proximity or even surrounded the ENS and outnumbered TIMP2^+^ Mφs, which were located further away from the ENS (Fig. 3E). To confirm that Cd206^+^ MhcII^hi^ Mφs have a neuroprotective on the ENS, we next aimed to assess their contribution on EGC proliferation *in vitro* (Fig. 3F). While Ly6c^+^ monocytes isolated 24h post-injury did not alter the proliferation of EGCs, MHCII^hi^ Mφs from 72h post-injury induced EGC proliferation, underscoring their role in supporting EGC function (Fig. 3G). These results establish that CCR2^+^ monocytes differentiate into two distinct Mφ subpopulations with pro-resolving and/or neurotrophic functions.

### CX3CR1-based mapping shows unique Mφ subsets similar to scRNAseq

To further define the phenotype of the Mφ subpopulations identified using scRNA at the protein level, we induced muscularis inflammation in CX3CR1^gfp/+^ mice (Fig. 4A-B). Upon muscularis inflammation, the percentage of homeostatic CX3CR1^hi^ Mφs was drastically reduced with a concomitant increase of CX3CR1^lo^ MHCII^hi^ and CX3CR1^lo^ MHCII^lo^ Mφs. As CCR2 and CD206 have been described as essential markers for recruited and anti-inflammatory Mφs respectively, their protein levels were determined in these CX3CR1^hi/lo^ Mφ subpopulations (Fig. 4C-H **and** Supplementary Fig. 4A-B). In line with previous findings, CX3CR1^hi^ MHCII^hi^ Mφs were mainly CD206^+^ and CCR2^-^ at homeostasis (Fig. 4C-E). However, this subpopulation was gradually replaced by a combination of CX3CR1^lo^ MHCII^lo^ and CX3CR1^lo^ MHCII^hi^ Mφs, resembling Timp2^+^ and Cd206^+^ Mφs respectively (Fig. 4C-H). Interestingly, CD206^+^ CX3CR1^lo^ MHCII^hi^ Mφs 10-fold outnumbered CX3CR1^lo^ MHCII^lo^ Mφs, further underscoring the importance of these anti-inflammatory Cd206^+^ MhcII^hi^ Mφs (Supplementary Fig. 4A-B). Overall, our data validated at protein level the presence of two distinct Mφ subpopulations during the resolution of muscularis inflammation which are derived from CCR2^+^ monocytes and are anti-inflammatory in nature.

**Figure 4:**
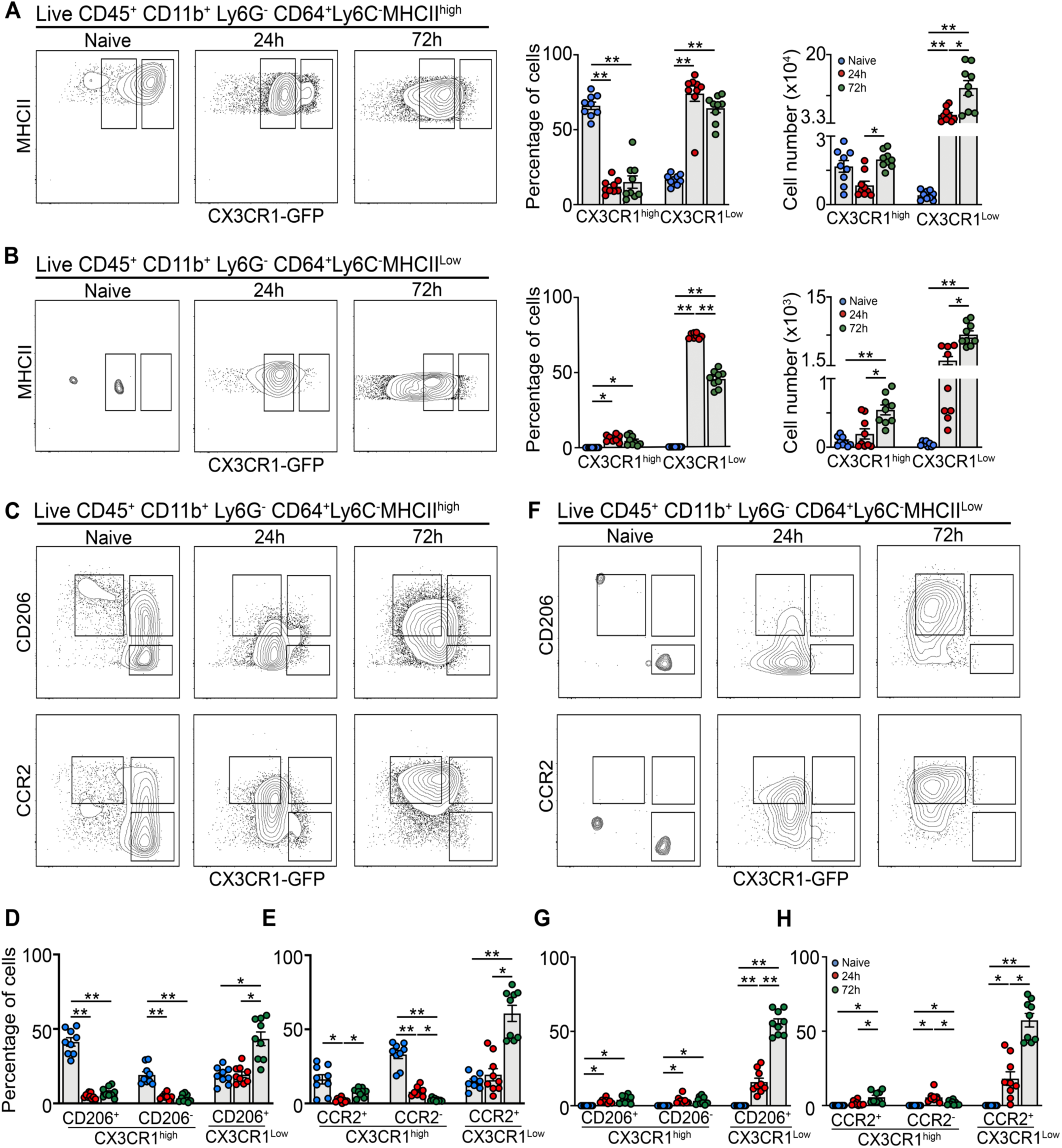
Flow cytometry validates unique Mφ subpopulations during muscularis inflammation. **A-H.** from naïve CX3CR1^GFP/+^ mice, 24h and 72h after muscularis inflammation**. A.** Contour plots representing CX3CR1 expression in Ly6C^-^ MHCII^hi^ Mφs (left). Percentages and absolute numbers of CX3CR1^hi^ and CX3CR1^lo^ cells from Ly6C^-^ MHCII^hi^ Mφs are shown as mean ± SEM (right). **B.** Contour plots representing CX3CR1 expression in Ly6C^-^ MHCII^lo^ Mφs (left). Percentages and absolute numbers of CX3CR1^hi^ and CX3CR1^lo^ cells from Ly6C^-^ MHCII^lo^ Mφs are shown as mean ± SEM (right). **C-H.** CD206 and CCR2 expression in Ly6C^-^ MHCII^hi^ and MHCII^lo^ Mφs. Contour plots representing CD206 or CCR2 expression in Ly6C^-^ MHCII^hi^ (C) Ly6C^-^ MHCII^lo^ Mφs (F). Percentages of CD206^+^ CX3CR1^hi^, CD206^-^ CX3CR1^hi^ and CD206^+^ CX3CR1^lo^ cells in the LY6C^-^ MHCII^hi^ Mφs (D) and MHCII^lo^ Mφs (G). Percentages of CCR2^+^ CX3CR1^hi^, CCR2^-^ CX3CR1^hi^ and CCR2^+^ CX3CR1^lo^ cells in the LY6C^-^ MHCII^hi^ Mφs (E) and MHCII^lo^ Mφs (H). One-way ANOVA; test *p<0.05; **p<0.01; ns=not significant.

Similar to our scRNAseq dataset, we observed a drastic reduction in the percentage of homeostatic CX3CR1^hi^ Mφs upon the induction of muscularis inflammation (Fig. 2C-D). To determine whether these CX3CR1^hi^ Mφs are replaced by incoming monocyte-derived Mφs, CX3CR1^+^ resident Mφs were mapped by using the tamoxifen-inducible CX3CR1^CreERT2^ strain backcrossed with Rosa26-LSL-YFP mice. By comparing circulating unlabeled blood monocytes with resident YFP^+^ Mφs, we could show that resident Mφs did not diminish in number during muscularis inflammation as quantified by immunofluorescence and flow cytometry (Supplementary Fig. 4C-F). Taken together, these results indicate that CX3CR1^hi^ Mφs were present alongside CX3CR1^lo^ mature Mφs during muscularis inflammation.

### EGC-derived CCL2 initiates the recruitment of monocytes upon muscularis inflammation

Monocytes originate from progenitors in the bone marrow (BM) and rely on local chemokine production such as MCP-1/CCL2 for their recruitment at the site of inflammation^30^. Considering the extensive interaction between CCR2^+^ monocytes and the ENS in the inflamed muscularis^6, 11^, we examined the production of the major chemoattractant Ccl2 in the muscularis during the early stage of surgery-induced inflammation. Analysis of isolated enteric ganglia 1.5h post-injury showed an increased gene expression of *Ccl2* and *Csf1* (Fig. 5A). Moreover, immunofluorescent images showed that CCL2 was specifically produced by GFAP^+^ EGCs (Fig. 5B), leading us to conclude that during the early stages of muscularis inflammation, EGCs likely initiate the recruitment of monocytes via CCL2 to stimulate tissue repair.

**Figure 5:**
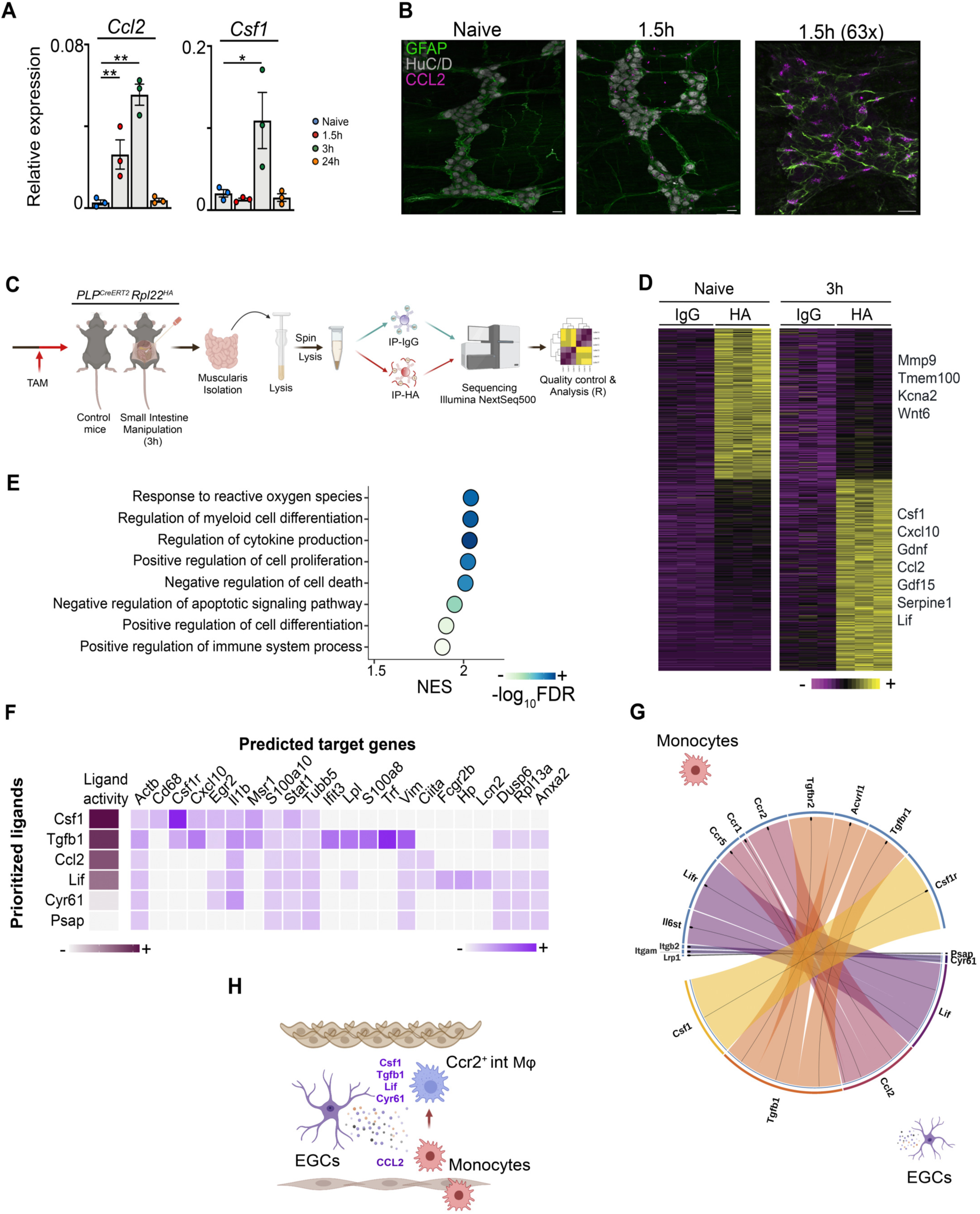
EGCs produce factors essential for the recruitment and differentiation of monocytes during inflammation. **A.** Relative mRNA levels for Ccl2 and Csf1 normalized to the housekeeping gene *rpl32* from ganglia isolated from the muscularis of the small intestine from naïve wild-type mice and 1.5h, 3h and 24h after muscularis inflammation. One-way ANOVA; test *p<0.05; **p<0.01; ns not significant. **B.** Immunofluorescent images of muscularis whole-mount preparations at homeostasis and 1.5h after the induction of muscularis inflammation stained for GFAP (green), HuC/D (grey) and CCL2 (purple). Scale bar (25x) 25 µm, (63x) 15 µm. **C.** Experimental overview of experiments in PLP-CreERT2 Rpl22^HA^ mice. **D.** Heatmap of differentially expressed genes between immunoprecipitated samples from naïve PLP-CreERT2 Rpl22^HA^ mice and 3h after intestinal manipulation. **E.** Selected significant GO terms enriched (GSEA) in PLP1^+^ EGCs 3h post-injury compared to naive PLP1^+^ EGCs. **F.** Heat map of ligand-target pairs showing regulatory potential scores between all positively correlated prioritized ligands and their target genes among the differentially expressed genes between Ly6c^+^ monocytes and Ccr2^+^ int Mφs. **G.** Circos plot showing top NicheNet ligand-receptor pairs between EGCs and Ly6c^+^ monocytes corresponding to the prioritized ligands in Fig. 5F. **H.** Schematic overview of interaction between EGCs and infiltrating Ly6c^+^ monocytes.

To further explore the role of EGCs during the early phase of muscularis inflammation, PLP^CreERT2^ Rpl22^HA^ mice were used, which allow immunoprecipitation (IP) of ribosome-bound mRNAs from PLP1^+^ EGCs to study the transcriptome of EGCs at homeostasis and during early inflammation (3h post-injury) (Fig. 5C **and** Supplementary Fig. 5A-B)^25, 31, 32^. Unlike pro-inflammatory reactive astrocytes during brain injury^33^, EGCs were activated early after muscularis inflammation and produced increased amounts of factors that are able to promote the differentiation of alternatively activated Mφs, such as *Lif, Serpine-1, Gdf15, Gdnf, Cxcl10* and *Csf1* (Fig. 5D). To explore which pathways are activated in EGCs, gene set enrichment analysis was performed. Positive normalized enrichment scores were identified for response to ROS and cytokines, indicating that EGCs are able to sense tissue damage in the muscularis microenvironment to initiate an appropriate immune response (Fig. 5E). Interestingly, there was also an overrepresentation of genes involved in the regulation of myeloid differentiation during early muscularis inflammation in EGCs (Fig. 5E **and** Supplementary Fig. 5C). These results suggest that EGCs might be involved in both the recruitment and differentiation of monocytes during the early stages of muscularis inflammation.

To determine whether EGCs are able to stimulate monocyte differentiation during muscularis inflammation, we analyzed the expression of ligand-receptor pairs based on differentially expressed genes in PLP1^+^ EGCs 3h post-injury and Ly6c^+^ monocytes undergoing differentiation into Ccr2^+^ int Mφs. Their expression data was combined with prior knowledge of signaling and gene regulatory networks using Nichenet^34^. To identify genes regulated by the identified ligands, putative ligand-gene interactions were scored by NicheNet according to their “regulatory potential” (Fig. 5F). We next assessed whether the expression of the receptors for the putative ligand-receptor interaction were altered when Ly6c^+^ monocytes underwent differentiation into Ccr2^+^ int Mφs (Supplementary Fig. 5D-E). By determining a threshold for the receptor expression, we identified the following ligand-receptor interactions between EGCs and differentiating monocytes: *Csf1-Csf1r, Tgfb1-Tgfbr2/Acvrl1/Tgfbr1, Ccl2-Ccr5/Ccr1/Ccr2, Lif-Lifr/Il6st, Cyr61-Itgam/Itgb2, and Psap-Lrp1* (Fig. 5G **and** Supplementary Fig. 5F). Taken together, EGCs are able to recruit monocytes to the site of inflammation, where they produce factors that have the potential to promote anti-inflammatory Mφ differentiation (Fig. 5H).

### EGC-derived CSF1 promotes the differentiation of monocytes into pro-resolving CD206^+^ Mφs

To directly assess the functional interaction between EGCs and monocytes, neurosphere-derived EGCs were analyzed by bulk RNAseq. The results revealed high expression of cytokines involved the induction of alternative Mφ differentiation (Supplementary Fig. 6A-C), similar to those observed *in vivo* in EGCs, including *Lif, Csf1, Tgfb1* and *Ccl2* (Fig. 5D, F). To demonstrate that EGC-derived factors are able to stimulate monocyte differentiation, an *ex vivo* model was set-up to co-culture BM monocytes with the supernatant of EGCs (Fig. 6A)^35^. Co-culture of BM monocytes with EGC supernatant resulted in an upregulation of several anti-inflammatory markers such as *Arg1, Il10* and *Mrc1*, while pro-inflammatory markers were downregulated, including *Il6* and *Il12* (Fig. 6B). In addition, when BM monocytes were exposed to bacterial components, a similar anti-inflammatory effect of EGC supernatant was observed compared to LPS alone (Supplementary Fig. 6D). Finally, to confirm that muscularis monocytes responded to EGC supernatant in a similar fashion as BM monocytes given their different origin, *ex vivo* experiments were performed on sorted Ly6C^+^ monocytes from the muscularis 24h post-injury (Fig. 6C). Also in this setting, monocytes acquired an anti-inflammatory phenotype upon co-culture with EGC supernatant similar to that seen in BM monocytes (Fig. 6D **and** Supplementary Fig. 6E). Furthermore, flow cytometric analysis of BM monocytes stimulated with EGC supernatant showed increased CD206 and reduced CCR2 expression, along with increased survival compared to control monocytes (Fig. 6E-H). Taken together, our *ex vivo* data suggest a direct interaction between EGCs and monocytes. As cellular interaction analysis by Nichenet identified Csf1-Csf1r as the top candidate for potential ligand-receptor interaction between EGCs and monocytes, we next aimed to determine whether EGC-derived CSF1 had the potential to induce anti-inflammatory CD206^+^ Mφs in our *ex vivo* co-culture model by antibody-mediated CSF1r blockade. Anti-CSF1r treatment in combination with EGC supernatant attenuated the differentiation of monocytes into pro-resolving Mφs with reduced CD206 and increased CCR2 expression compared to cells stimulated with EGC supernatant (Fig. 6F **and** Supplementary Fig. 6F-H). Furthermore, we observed increased cell death of monocytes in EGC supernatant + anti-CSF1r as compared to EGC control supernatant at 48h (Fig. 6E **and** Supplementary Fig. 6G). Altogether, these data provide evidence for a direct interaction between monocytes and EGC-derived ligands *ex vivo* which stimulates the differentiation into pro-resolving CD206^+^ Mφs, which are essential for the resolution of muscularis inflammation.

**Figure 6:**
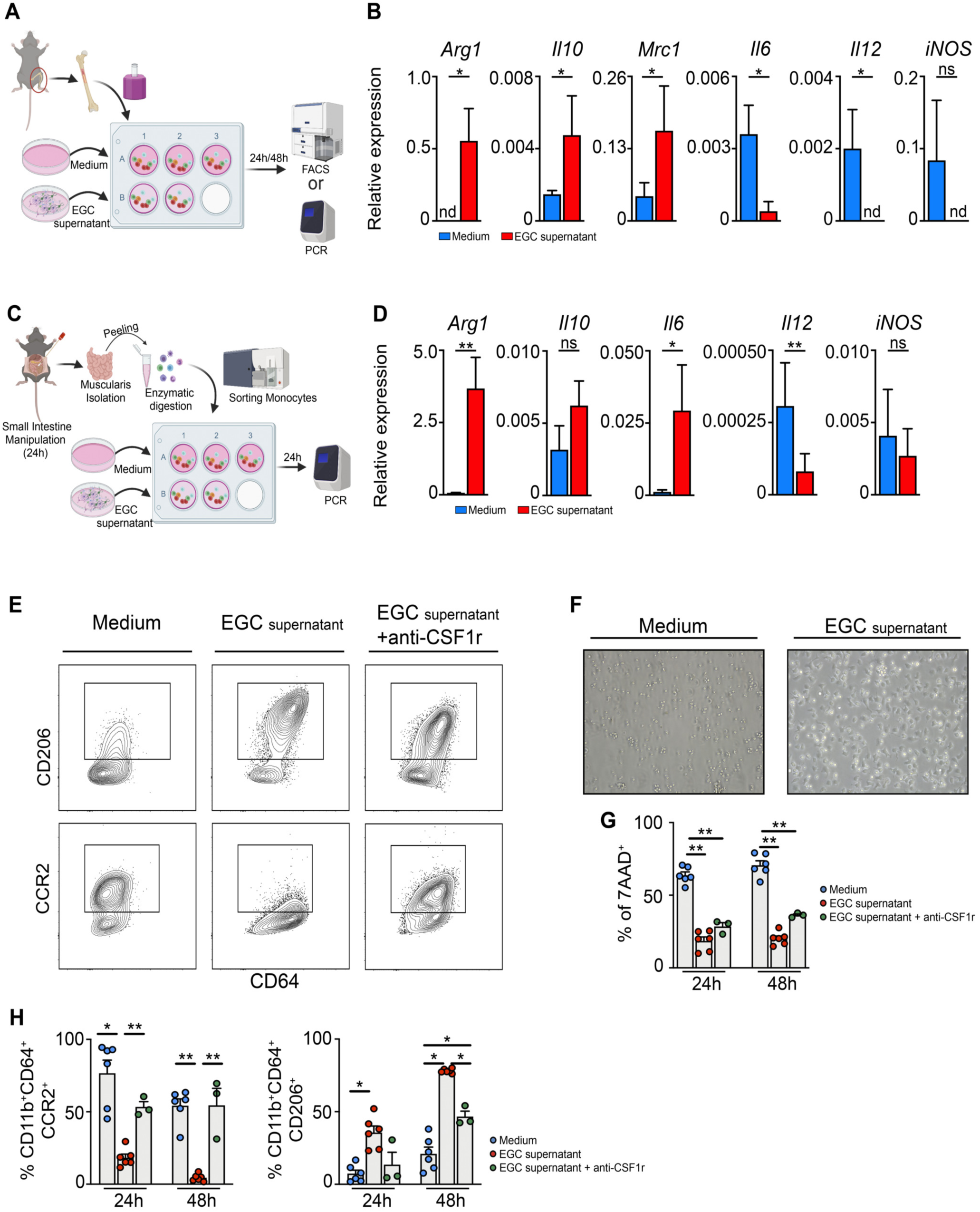
Enteric glia-derived CSF1 stimulates the differentiation of monocytes into pro-resolving CD206^+^ Mφs. **A.** Experimental outline of *in vitro* BM monocytes stimulated with supernatant of EGCs. **B.** Relative mRNA levels for pro- and anti-inflammatory cytokines normalized to the housekeeping gene *rpl32* in BM monocytes cultured for 24h with or without EGC supernatant. **C.** Experimental outline of *in vitro* experiment using sorted Ly6C^+^ MHCII^-^ monocytes stimulated with EGC supernatant for 24h. **D.** Relative mRNA levels for of pro- and anti-inflammatory mediators normalized to the housekeeping gene *rpl32* in sorted Ly6C^+^ MHCII^-^ monocytes stimulated with or without EGC supernatant. **E-H**. BM monocytes were cultured for 24-48h with/without EGC supernatant and/or supplemented with anti-CSF1r antibody. **E**. Contour plots of BM monocytes showing expression of CD206 (top) or CCR2 (bottom) upon culture for 48h with/without EGC supernatant and supplemented with anti-CSF1r antibody. **F**. Brightfield images of monocytes upon culture for 24h with or without EGC supernatant. **G**. Quantification of 7-AAD^+^ cells in BM monocytes cultured for 24h-48h with/without EGC supernatant or supplemented with anti-CSF1r antibody. **H**. Percentages of CD11b^+^ CD64^+^ cells in live CD45^+^ population, and CCR2^+^ and CD206^+^ cells in live CD45^+^ CD11b^+^ Ly6G^-^ CD64^+^ population. A-D. T-test. G-H. one-way ANOVA. *p<0.05; **p<0.01.

## Discussion

Previous studies have shown that monocyte-derived macrophages are essential to resolve inflammation and restore gut motility following muscularis inflammation^11, 28^. To date, it remains to be determined which Mφ subpopulations are responsible for the recovery following muscularis inflammation and which factors induce the pro-resolving Mφ phenotype essential for tissue repair. Our work provides a comprehensive overview of the differentiation of incoming monocytes into pro-resolving Mφs during muscularis inflammation, thereby characterizing gene expression dynamics during monocyte-to-Mφ transitions and uncovering transcriptional regulators of different Mφ subpopulations. Furthermore, we have shown that upon muscularis injury the gene expression pattern of EGCs changes dramatically leading to an increased production of the monocyte chemoattract CCL2 and elevated levels of multiple mediators that could be involved in the differentiation of monocytes towards anti-inflammatory Mφs, such as CSF1, during this paradigm.

Using time-resolved scRNAseq of CD45^+^ immune cells from the inflamed muscularis, we identified different monocyte and Mφ subpopulations at homeostasis and during acute inflammation and the resolution of inflammation. During acute muscularis inflammation, Ly6c^+^ monocytes were present along with other transient monocyte/Mφ subpopulations such as Fabp5^+^ and Arg1^+^ mono/Mφ, showing substantial heterogeneity in monocytic cells during acute inflammation, rather than a single homogenous wave of Ly6c^+^ monocytes. Additionally, we identified Ccr2^+^ int Mφs as a transitional state between monocytes and Cd206^+^ Mφs. These intermediate cells were mainly present near the branching point or in the branch of the trajectory diverging towards Cd206^+^ Mφs and resembled Ly6C^+^ MHCII^+^ immature Mφs characterized by flow cytometry. Furthermore, we identified two main pro-resolving Mφ subpopulations during the resolution of muscularis inflammation characterized by Cd206^+^ MhcII^hi^ and Timp2^+^ MhcII^lo^ gene signatures, which are transcriptionally distinct from homeostatic Cx3cr1^+^ Mφs. Trajectory interference analysis during muscularis inflammation showed that these subpopulations are derived from infiltrating Ly6c^+^ monocytes. In line, scRNAseq analysis showed that these two myeloid subpopulations are absent from the inflamed muscularis of CCR2 deficient mice, further supporting their monocytic origin. Accordingly, flow cytometric analysis of CX3CR1^GFP/+^ mice during muscularis inflammation showed that CX3CR1^lo^ MHCII^hi^ and CX3CR1^lo^ MHCII^lo^ Mφs are derived from CCR2^+^ monocytes and resemble Cd206^+^ and Timp2^+^ Mφs, respectively. Both phenotypically and functionally, Cd206^+^ MhcII^hi^ Mφs showed high resemblance to homeostatic Cx3cr1^+^ Mφs with a similar gene signature and activity of specific regulons (e.g. *Junb, Irf8)*^36, 37^, while Timp2^+^ MhcII^lo^ Mφs displayed an alternative gene signature and regulon activity (e.g. *Cebpb, Gata6)*^38, 39^. In this regard, Cd206^+^ Mφs were shown to have high regulon activity of *Irf8*, which regulates the expression of genes important for the activation of antigen-presenting cells and T cells (*Cd86* and *Cd40*) and for lymphoid and myeloid chemotaxis (*Cxcl9, Cxcl10, Ccl4, Ccl8* and *Ccl12*)^37^. In addition, these cells upregulated genes involved in efferocytosis and exhibited high *MhcII* gene expression. These data indicate that CD206^+^ Mφs might respond to apoptotic cells in their vicinity and undergo alternative activation in response to efferocytosis. On the other hand, Timp2^+^ Mφs have high regulon activity of *Cebpb*, which controls the expression of anti-inflammatory genes such as *Msr1, Il10, II13ra*, and *Arg1*. These cells seem to have a high metabolic activity as demonstrated by their GO terms. Studies in small and large peritoneal Mφs could be further instrumental in understanding Mφ biology during muscularis inflammation, as these cells resemble Cd206^+^ and Timp2^+^ Mφs, respectively, based on gene expression and regulon activity^40, 41^. Furthermore, we also found other subpopulations that were characterized by highly specific gene signatures and regulon activities, such as Fabp5^+^ Mφs with enhanced *Nr1h3 (LXRα)* and *Pparγ* regulon activity, which may be involved in iron handling (*Flt1, Fth1, Hmox1*) and detoxification (*Mt1, Akr1a1, Prdx1, Cd36*). Further studies will be essential to unravel the functional roles of different Mφ subsets during muscularis inflammation by defining their precise tissue localization and assessing their contribution to the pathophysiological response during local tissue damage.

The muscularis microenvironment is mainly regulated by the ENS, which is composed of a network of enteric neurons and EGCs. Here, Mφs are essential to support neuronal function via the secretion of BMP-2^1, 6^, while neurons in turn produce CSF1, vital for Mφ development^6^. In addition, CSF1 production has also recently been identified in EGCs^19^. This close neuro-immune interaction between the tissue microenvironment and Mφs at homeostasis and during disease is also found in other neuronal tissue niches. For instance, in the central nervous system, there is extensive crosstalk between astrocytes and microglia, critical for brain homeostasis^42^. Upon brain injury, microglia are the first cells to be recruited to the site of damage to phagocytose dead cells and debris^43^, similar to what is observed during muscularis inflammation^11, 28^. Concurrently, reactive astrocytes are activated to release pro-inflammatory mediators that affect microglia and can recruit different leukocytes. Subsequently, they increase their expression of GFAP and undergo morphological changes leading to the formation of glial scars^44^. Similarly, we have shown a comparable process taking place at the level of the ENS, in which damage-activated EGCs closely interact with Mφs in the muscularis and act as safeguards of the microenvironment by initiating an inflammatory response via CCL2 to protect the tissue from further damage. As opposed to the detrimental effects of astrocytes in brain injury, EGCs induce monocyte differentiation towards pro-resolving CD206^+^ Mφs by the production of CSF1, which seems to be important for the resolution of ENS damage as Mφs are able to stimulate EGC proliferation. Similarly, upon Mφ depletion, red pulp fibroblasts transiently produce the monocyte chemoattractants CSF1, CCL2 and CCL7, thereby contributing to monocyte replenishment^45^. Recently, it was also shown that during experimental autoimmune encephalomyelitis, astrocytes provide soluble signals to pro-inflammatory Mφs to induce an anti-inflammatory phenotype^46^.

Overall, these insights may give rise to novel therapeutic approaches to treat patients affected by intestinal inflammatory disorders to avoid complications related to alterations of the ENS and reduced GI motility. Our findings on the impact of anti-inflammatory signals from EGCs on myeloid cells could be extrapolated beyond the context of the intestinal inflammation and may be valuable in the context of several other acute and chronic inflammatory conditions and autoimmune diseases.

## Acknowledgments

The authors would like to thank Iris Appeltans, Naomi Fabre, Tine Gommers and Karlien Vranken (TARGID, KU Leuven) for their technical assistance, and Pier Andrée Penttila and Reena Chinnaraj (FACS Core, KU Leuven) for their assistance with flow cytometry and sorting, the GIGA-Genomics platform (University of Liège) for their assistance in our scRNAseq experiments. Images were recorded at the Cell and Tissue Imaging Cluster (KU Leuven) using a Zeiss LSM 880 – Airyscan (supported by Hercules AKUL/15/37_GOH1816N and FWO G.0929.15 to Pieter Vanden Berghe) and a Zeiss LSM 780 – SP Mai Tai HP DS (supported by Hercules AKUL/11/37 and FWO G.0929.15 to Pieter Vanden Berghe). BioRender was used for making graphical images.

## Author contributions

G.M. and S.I. conceptualized this study; S.I. and M.S. designed the experiments; M.S., S.A., V.D.S., G.G., N.S. L.V.B., Q.W., D.P., J.K., L.M., I.P. and S.T. performed the experiments; analyzed data; M.S., S.A., S.I. and G.M. interpreted data; L.B., J.V.G., J.T., S.J. and T.M. provided vital reagents; M.H., P.V., J.V.G., G.B., J.T., S.J. and T.M. gave technical support and conceptual advice, M.S., S.A., S.I. and G.M. contributed to manuscript writing. All authors edited and revised the manuscript.

## Supplementary information

### Supplementary methods

#### Immunofluorescence

The small intestine was removed and flushed with ice-cold PBS to remove luminal contents. Then, the intestine was cut open longitudinally before stretching and was fixed for 30 min in 4% PFA at room temperature. Next, the tissue was washed three times with PBS and the mucosa and submucosa were removed with a forceps to obtain whole-mounts of the muscularis, after which it was permeabilized and blocked in 0.3% Triton X-100 and 3% BSA in PBS for 2h at room temperature. Subsequently, samples were incubated for 48h at 4°C with the following primary antibodies: chicken anti-neurofilament (Abcam; ab72996), rat anti-GFP (Nacalai Tesque; GF090R), goat anti-CCL2 (R&D Systems; AF-479-NA), rabbit anti-GFAP (Dako; Z0334), chicken anti-GFAP (Abcam; ab4674), goat anti-TIMP2 (R&D systems, AF971), rabbit anti-CD206 (Abcam, ab64693), rabbit anti-HA (CST; Rb#3724), human ANNA-1 serum anti-HuC/D (kindly provided by Prof. Lennon V. A. (Mayo Clinic, Rochester, Minnesota, USA)^1^. Afterwards, samples were washed in PBS and incubated with DAPI (4′,6-Diamidine-2′-phenylindole dihydrochloride; Sigma-Aldrich) combined with the secondary antibodies: donkey anti-chicken Cy5 (Jackson), donkey anti-rat Alexa Fluor 488 (Invitrogen), donkey anti-rabbit Cy3 (Jackson), donkey anti-human Cy5 (Jackson) and donkey anti-rabbit FITC. All primary and secondary antibodies were dissolved in 0.3% Triton X-100 and 3% BSA in PBS. Finally, samples were rinsed three times in PBS and mounted with SlowFade Diamond Antifade mountant (Invitrogen). Images were recorded on a LSM780 or LSM880 multiphoton microscope (Zeiss) and a 25x or 63x objective (Zeiss) was used. Identical settings were used for all conditions in one experiment. Images were processed and analyzed using ImageJ (NIH) or Imaris software (Bitplane).

#### Lineage Tracing of CX3CR1^+^ Macrophages

To induce CreERT2 recombinase activity to trace CX3CR1^+^ muscularis Mφs, Cx3cr1^CreERT2^ mice were crossed with Rosa26-LSL-YFP mice to obtain double heterozygous mice. Mice aged 8 weeks were injected three times subcutaneously with 4 mg TAM (Sigma-Aldrich, St. Louis, USA) per 30 g body weight dissolved in corn oil (Sigma-Aldrich) every other day as previously described^2^. Mice were sacrificed 4 weeks after the first tamoxifen injection.

#### Isolation and digestion of the muscularis

The small intestine was removed and flushed with ice-cold PBS. The muscularis was carefully removed with a forceps, cut in small pieces with scissors and digested in 2 mg/mL collagenase type IV (Gibco) in RPMI-1640 (Lonza) supplemented with 2% HEPES (Gibco), 2% FBS and 5U/mL DNase I (Roche) for 30 min at 37°C with continuous stirring. The resulting cell suspension was blocked using FACS buffer and passed through a 70 μm cell strainer (BD Falcon), after which cells were spun down at 400 x g for 8 min at 4°C. Cells were counted using the Countess II FL (Thermo Fisher).

#### Flow Cytometry and Sorting of Live Cells

Single cell suspensions were blocked with rat anti-mouse CD16/CD32 (Fc block, BD Biosciences) for 10 min and afterward incubated with fluorophore-conjugated anti-mouse antibodies at recommended dilutions for 20 min at 4°C. Dead cells were excluded using 7-AAD (BD Biosciences) or Fixable Viability Dye eFluor 450 (eBiosciences). During FACS acquisition, doublets were excluded. FACS antibodies used can be found in Table 1. Samples were acquired using a Symphony (BD Biosciences) and analyzed with FlowJo software (version 4.6.2, Treestar). For cell sorting, a BD Aria III (BD Biosciences) was used.

**Table 1:** FACS antibodies

| Anti | Conjugate | Company | Cat. No. | Clone |
| --- | --- | --- | --- | --- |
| CD45 | APC-eFluor 780 | eBioscience | 47-0451-82 | 30-F11 |
| CD45 | BV510 | Biolegend | 103138 | 30-F11 |
| CD45 | FITC | BD Pharmingen | 553772 | 104 |
| CD11b | PE-Cy7 | BD Pharmingen | 552850 | M1/70 |
| Ly6C | BV421 | BD Pharmingen | 562727 | AL-21 |
| Ly6C | PE | BD Pharmingen | 560592 | AL-21 |
| CD64a and b | BV711 | Biolegend | 139311 | X54-5/7.1 |
| CD64a and b | BV421 | Biolegend | 139309 | X54-5/7.1 |
| CD206 | Alexa Fluor 647 | BioLegend | 141712 | C068C2 |
| CCR2 | BUV395 | BD Pharmingen | 747972 | 475301 |
| Ly6G | BUV563 | BD Pharmingen | 612921 | IA8 |
| Ly6G | APC | BD Pharmingen | 560599 | IA8 |
| MHCII (I-A/I-E) | APC-eFluor 780 | eBioscience | 47-5321-82 | M5/114.15.2 |

#### Muscularis ganglia isolation from adult mice

The small intestine was flushed with cold bicarbonate/CO_2_ KREBS buffer. Thereafter, the myenteric plexus was collected by removing the mucosa and submucosa as previously described^3^. The extracted tissue was subsequently placed in GentleMACS™ tubes (Miltenyi Biotec, Germany) with 5mL of pre-warmed dissociation medium DMEM/F12 (Lonza, Belgium) supplemented with 10% FBS (Biowest, USA), 100U/mL Penicillin/Streptomycin, 2mM L-glutamine (Lonza, Belgium), 45mg/mL NaHCO3, 0.01 mg/mL Gentamicin and 0.25 µg/mL Amphotericin B (Thermo Fisher). Additionally, 400µl BSA (50g/L), 250µl protease (20mg/mL) and 250µl collagenase type IV (20mg/mL) (Sigma-Aldrich, Belgium) were added for enzymatic dissociation of tissue. The GentleMACs™ tube was then placed into the GentleMACS™ Dissociator (Miltenyi Biotec, Germany) for mechanical dissociation for 2 min at room temperature. The tube was subsequently placed in a water bath at 37°C for 13 min with shaking. Subsequently, a second mechanical dissociation was performed, followed by stopping the enzymatic dissociation by addition of cold KREBS buffer. Samples were spun down at 407 x g at 4°C and re-suspended in dissociation medium without enzymes, plated in a petri dish and placed under an inverted binocular microscope, where the ganglia were picked up with a pipette tip and were lysed with RLT buffer for assessment of RNA expression levels.

#### Neurosphere-derived embryonic enteric glial cell culture

Neurosphere-derived EGCs were obtained as previously described^4^. Briefly, total intestines from E14.5 C57BL/6J mice were digested with collagenase D (0.5 mg/mL; Roche) and DNase I (0.1mg/mL; Roche) in DMEM/F-12, GlutaMAX, supplemented with 1% HEPES, streptomycin/penicillin and 0.1% β-mercaptoethanol (Gibco) for approximately 30 min at 37°C under gentle agitation. Cells were cultured for 1 week in a CO_2_ incubator at 37°C in DMEM/F-12, GlutaMAX, streptomycin and penicillin and 0.1% β-mercaptoethanol (Gibco) supplemented with B27 (Gibco), 40 ng/mL EGF (Gibco) and 20 ng/mL FGF (Gibco). After 1 week of culture, neurospheres were treated with 0.05% trypsin (Gibco), transferred into PDL (Sigma-Aldrich) coated plates and differentiated in DMEM supplemented with 10% FBS, 1% HEPES, glutamine, streptomycin and penicillin and 0.1% β-mercaptoethanol (Gibco) until confluence for 5 days to obtain EGC supernatant.

#### RNA sequencing of neurosphere-derived EGC

RNA was isolated using RNeasy Micro Kit according to the manufacturer’s instructions. A cDNA indexed library was prepared using the Illumina TruSeq RNA library Kit and sequencing was performed on the Illumina NextSeq500 platform at the Nucleomics core (VIB, Leuven). The reads were preprocessed by trimming for quality and to remove adapters (cutadapt 1.15 and FastX 0.0.14) followed by filtering for quality (FastX 0.0.14 and ShortRead 1.40.0) and removal of contaminants (bowtie 2.3.3.1). The reads were then mapped against the reference genome GRCm3882 using STAR (2.5.2b). Further filtering, sorting and indexing was done using samtools 1.5 and counts file was prepared and differential expression analysis was performed using EdgeR.

#### Monocyte isolation

Mouse BM monocytes were isolated from WT mice. Briefly, the tibia and femur of mice were dissected. BM cells were flushed with DMEM high glucose (Lonza) supplemented with 10% FBS. After cells were collected and counted, monocytes were isolated with the EasySep™ Mouse Monocyte Isolation Kit (Stem Cell Technologies, Vancouver, Canada) according to the manufacturer’s instructions. Next, 10^5^ monocytes were stimulated for 24-48h with EGC supernatant, CSF-1 (50 ng/mL; PeproTech) and/or anti-CSF1r (5µg/mL; BE0213; BioXCell) at 37°C. Cells were stimulated with LPS (100 ng/ml; Sigma-Aldrich) 6 hours prior to the end of the experiment. Monocytes were analyzed by flow cytometry or lysed with RLT buffer.

#### EGC proliferation assay

After 1 week of culture, enteric neurospheres were collected and dissociated to generate a single-cell suspension by using the NeuroCult™ Chemical Dissociation Kit (Mouse) (Stem Cell Technologies, CatN°05707). Next, cells were filtered through a 70µm cell strainer and counted using a burker chamber. EGCs were transferred into poly-D-Lysine (Gibco, CatN°A3890401) coated 96-well plates at a concentration of 5000 cells per well and 100 µl of conditioned medium was added to each well. To obtain conditioned medium from myeloid cells, Ly6C^hi^ monocytes from 24h post-injury and Ly6C^-^ MHCII^hi^ Mφs from 72h post-injury were sorted from the muscularis and cultured at a concentration of 50.000 cells/well per 400µl medium with 0.5% FBS during 24h. After 5 days of culture in (un)conditioned media, the proliferation of the EGCs was measured by using the Cell Counting Kit-8 (Tebu-bio, CatN°CK04-05). After 2 hours of incubation with the CCK-8 solution, the absorbance was measured at 450nm and 600nm (background signal) using a microplate reader. Data were expressed as induction with respect to EGCs cultured with unconditioned medium.

#### RNA extraction and gene expression of ganglia and myeloid cells

Picked ganglia from the small intestine and monocytes/macropahges were lysed in RLT buffer and stored at −80°C. RNA extraction was performed using RNeasy Mini Kit for tissue and high cell numbers (Qiagen) or RNeasy Micro Kit for low cell numbers following manufacturer’s instructions. Total RNA was transcribed into cDNA by the High-Capacity cDNA Reverse Transcription Kit (Thermo Fisher) according to manufacturer’s instructions. Quantitative real-time transcription polymerase chain reactions (RT-PCR) were performed with the LightCycler 480 SYBR Green I Master (Roche) on the Light Cycler 480 (Roche). Results were quantified using the 2-ΔΔCT method^5^. Expression levels of the genes of interest were normalized to the expression levels of the reference gene Rpl32. Results are expressed as mean values and primer sequences used are listed in Table 2.

**Table 2:**
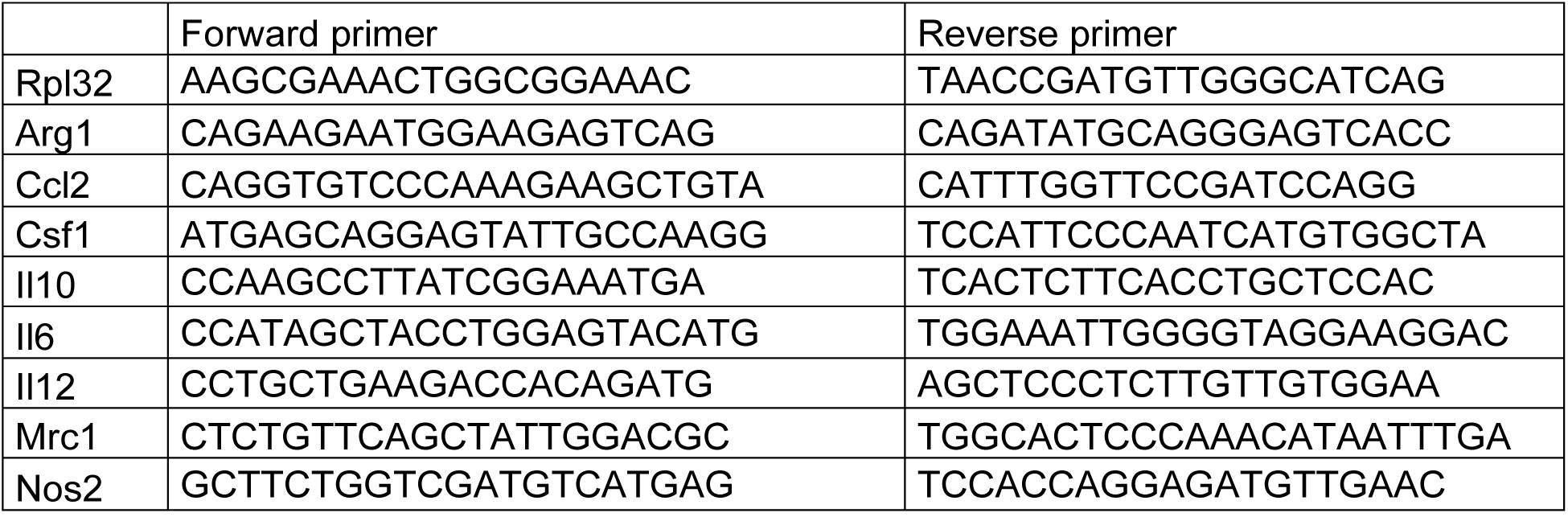
List of RT-PCR primers

#### Isolation and sequencing of ribosome-associated mRNA from EGCs

To induce gene recombination of Rpl22^HA^ in PLP1^CreERT2^ Rpl22^HA^ transgenic mice at 10-12 weeks of age, mice were injected twice intraperitoneally with 100 µl of 15 mg/mL tamoxifen (Sigma-Aldrich) dissolved in Miglyol 812 (Caesar & Loretz, Hilden, Germany). All animals were sacrificed 3-4 weeks after the first injection. The whole small intestine of each mouse was peeled by removing the mucosa and submucosa and samples were flash-frozen in liquid nitrogen and stored at −80°C until further processing. Samples were immunoprecipitated at the Weizman Instiute (Israel) according to the published protocol^6, 7^. In brief, samples were homogenized in 4 mL cold homogenization buffer (50 mM Tris, pH 7.4, 100 mM KCl, 12 mM MgCl2, 1% NP-40, 1 mM DTT, 1:100 protease inhibitor (Sigma-Aldrich), 200 U/mL RNasin (Promega) and 0.1 mg/mL cycloheximide (Sigma-Aldrich)) with a Dounce homogenizer (Sigma-Aldrich) until the suspension was homogeneous. To remove cell debris, 2 mL of the homogenate was transferred to a microcentrifuge tube and centrifuged at 10,000 g at 4°C for 10 min. Supernatants were transferred and split, and 8 μL (10 µg) of anti-HA antibody (H9658, Sigma-Aldrich) or 20 μL (10 µg) of mouse monoclonal IgG1 antibody (Sigma-Aldrich, Cat# M5284) was added to the supernatant, followed by 4h of incubation with slow rotation at 4°C. Meanwhile, Dynabeads Protein G (Thermo Fisher), were equilibrated to homogenization buffer by washing three times. At the end of 4 h of incubation with antibody, beads were added to each sample, followed by incubation overnight at 4°C. Next, samples were washed three times with high-salt buffer (50 mM Tris, 300 mM KCl, 12 mM MgCl2, 1% NP-40, 1 mM DTT, 1:200 protease inhibitor, 100 U/mL RNasin and 0.1 mg/mL cycloheximide). At the end of the washes, beads were magnetized and excess buffer was removed, 150 µl lysis buffer was added to the beads and RNA was extracted with Dynabeads mRNA Direct purification kit (Thermo Fisher) according to manufacturer’s instructions. RNA was eluted in 6 μl H_2_O and used for RNA-seq.

#### RNAseq library preparation and analysis of immunoprecipitated samples

Immunoprecipitated samples were processed and analyzed according to published protocols^6, 7^. In brief, mRNA was captured with Dynabeads oligo(dT) (Life Technologies) according to manufacturer’s guidelines. A bulk variation of MARSseq^8^ was used to prepare libraries for RNA-seq. RNA was reversed transcribed with MARSseq barcoded RT primer in a 10 mL volume with the Affinity Script kit (Agilent). Reverse transcription was analyzed by qRT-PCR and samples with a similar CT were pooled (up to eight samples per pool). Each pool was treated with Exonuclease I (NEB) for 30 min at 37°C and subsequently cleaned by 1.2 x volumes of SPRI beads (Beckman Coulter). Subsequently, the cDNA was converted to double-stranded DNA with a second strand synthesis kit (NEB) in a 20mL reaction, incubating for 2 h at 16°C. The product was purified with 1.4 x volumes of SPRI beads, eluted in 8 mL and in vitro transcribed (with the beads) at 37°C overnight for linear amplification using the T7 High Yield RNA polymerase in vitro transcription kit (NEB). Following in vitro transcription, the DNA template was removed with Turbo DNase I (Ambion) for 15 min at 37°C and the amplified RNA (aRNA) was purified with 1.2 volumes of SPRI beads. The aRNA was fragmented by incubating in Zn^2+^ RNA fragmentation reagents (Ambion) for 3 min at 70°C and purified with 2 x volumes of SPRI beads. The aRNA was ligated to the MARS-seq ligation adaptor with T4 RNA Ligase I (NEB). The reaction was incubated for 2 h at 22°C. After 1.5 x SPRI cleanup, the ligated product was reverse transcribed using Affinity Script RT enzyme (Agilent) and a primer complementary to the ligated adaptor. The reaction was incubated for 2 min at 42°C, for 45 min at 50°C, and for 5 min at 85°C. The cDNA was purified with 1.5 x volumes of SPRI beads. The library was completed and amplified through a nested PCR reaction with 0.5mM of P5_Rd1 and P7_Rd2 primers and PCR ready mix (Kappa Biosystems). The amplified pooled library was purified with 0.7 x volumes of SPRI beads to remove primer leftovers. Library concentration was measured with a Qubit fluorometer (Life Technologies) and mean molecule size was determined with a 2200 TapeStation instrument. RNaseq libraries were sequenced using the Illumina NextSeq 2000.

Data analysis was performed by using the UTAP transcriptome analysis pipeline^9^. Raw reads were trimmed using cutadapt with the parameters: -a AGATCGGAAGAGCACACGTCTGAACTCCAGTCAC -a ‘‘A–times 2 -u 3 -u -3 -q 20 -m 25). Reads were mapped to the genome (mm10, Gencode annotation version 10.0) using STAR (v2.4.2a) with the parameters –alignEndsType EndToEnd, –outFilterMismatchNoverLmax 0.05, –twopassMode Basic, –alignSoftClipAtReferenceEnds No. The pipeline quantifies the 3’ of Gencode annotated genes. UMI counting was done after marking duplicates (in-house script) using HTSeq-count in union mode. Only reads with unique mapping were considered for further analysis, and genes having minimum 10 reads in at least one sample were considered. Gene expression levels were calculated and normalized using DESeq2 (1.26.0) with default parameters except alpha = 0.05^10^.

#### Single cell RNA sequencing – Library preparation and sequencing

Single cell RNA sequencing libraries were prepared using the 10X Genomics platform according to manufacturer’s instructions (Single cell 3’ solution) and sequencing was performed on Illumina NextSeq500 (Illumina). Cell Ranger software (v3.0.2; 10X Genomics) was used to demultiplex Illumina BCL files to FASTQ files (cellranger mkfastq) for alignment to mouse GRCm38/mm10 genome, filtering, UMI counting, and to produce gene-barcode matrices. Library preparation, sequencing and sample demultiplexing were performed at the Genomics Platform of the GIGA Institute (Liège University, Belgium).

#### Single-cell RNA sequencing Clustering and Dimensionality reduction

The filtered count matrices obtained after pre-processing with Cellranger were concatenated to obtain a combined raw count matrix which was then analyzed using the Seurat R package (3.1.3). Cells with less than 200 genes, cells with more than 10% mitochondrial genes and genes with expression in less than 3 cells were discarded from the analysis. SCTransform function was used for normalization and scaling. A regression of the percentage of mitochondrial genes was performed during the SCTransform step. Next, principal component analysis was used to reduce the dimension of the data before UMAP construction and clustering. The cells annotated as monocytes and Mφs were then employed to subset original concatenated counts matrix. The subset matrix was then subjected to the same process of normalization, scaling (SCTransform), principal component analysis, UMAP projection and clustering. The MHC II signature score was calculated using ‘AUCell’ R package as the ‘area under the curve’ with the following genes - H2-Ab1, H2-Aa, H2-Eb1, H2-Eb2, H2-DMa, H2-DMb1, H2-DMb2. During the re-clustering of monocytes and Mφs from WT and CCR2^-/-^ mice, 67 cells appeared to be granulocytes/DCs (based on *Cxcr2* and *Cd209a* expression) and were not included in the re-clustering analysis of mono/Mφs. Analysis plots were generated using the R packages ggplot (3.2.1), pheatmap (1.0.12) and EnhancedVolcano (1.4.0). Markers for different clusters were determined using a Wilcoxon rank test with FindMarkers and FindAllMarkers functions in Seurat. Non-default parameters used at each step during clustering or re-clustering with Seurat are given in Table 3.

**Supplementary Table 3:**
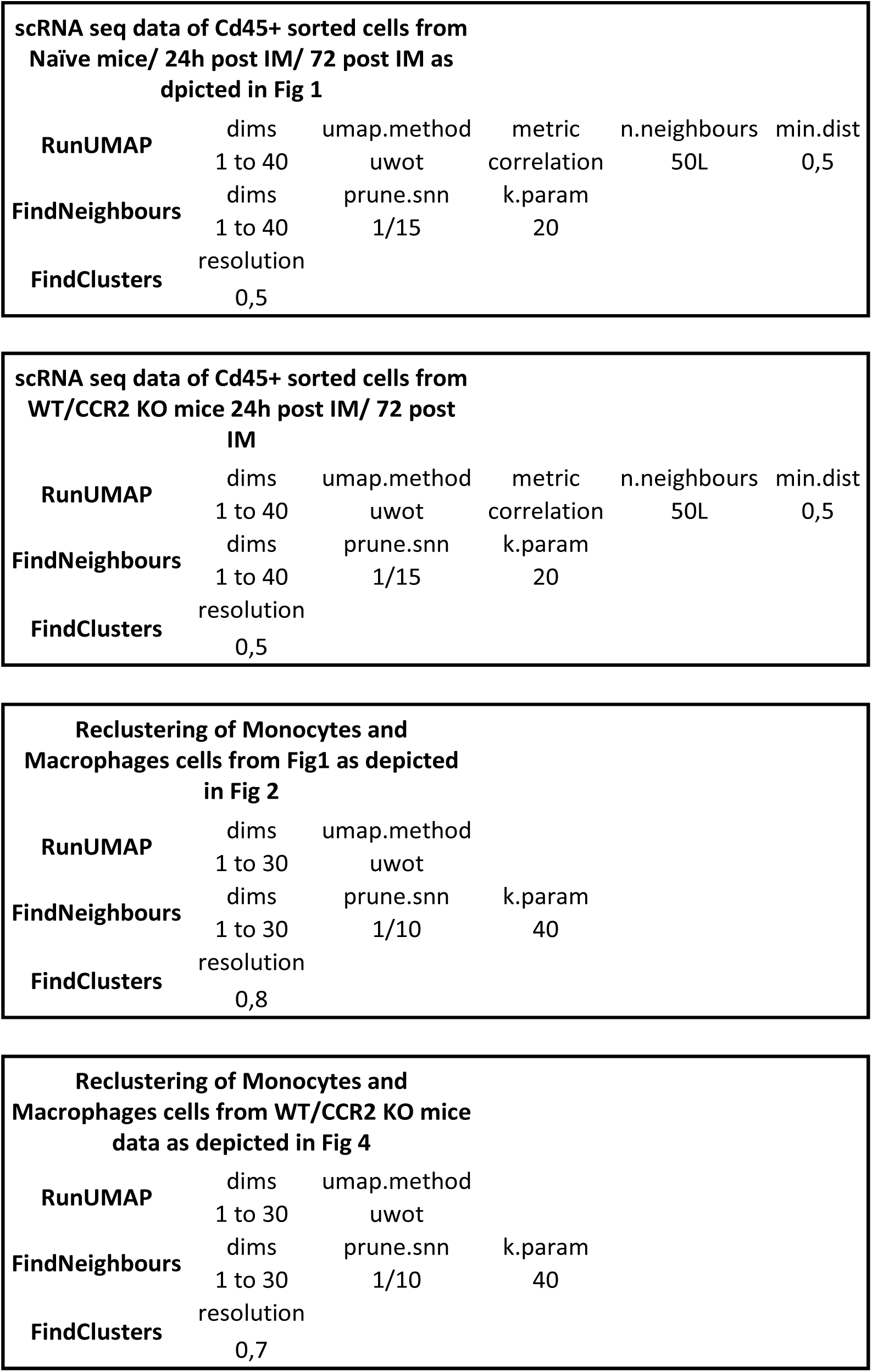
Non-default parameters Seurat.

| scRNA seq data of Cd45+ sorted cells from<br>Naïve mice/ 24h post IM/ 72 post IM as<br>depicted in Fig 1 |  |  |  |  |  |
| --- | --- | --- | --- | --- | --- |
| **RunUMAP** | dims | umap.method | metric | n.neighbours | min.dist |
|  | 1 to 40 | uwot | correlation | 50L | 0,5 |
| **FindNeighbours** | dims | prune.snn | k.param |  |  |
|  | 1 to 40 | 1/15 | 20 |  |  |
| **FindClusters** | resolution |  |  |  |  |
|  | 0,5 |  |  |  |  |

| scRNA seq data of Cd45+ sorted cells from<br>WT/CCR2 KO mice 24h post IM/ 72 post<br>IM |  |  |  |  |  |
| --- | --- | --- | --- | --- | --- |
| **RunUMAP** | dims | umap.method | metric | n.neighbours | min.dist |
|  | 1 to 40 | uwot | correlation | 50L | 0,5 |
| **FindNeighbours** | dims | prune.snn | k.param |  |  |
|  | 1 to 40 | 1/15 | 20 |  |  |
| **FindClusters** | resolution |  |  |  |  |
|  | 0,5 |  |  |  |  |

| Reclustering of Monocytes and<br>Macrophages cells from Fig1 as depicted<br>in Fig 2 |  |  |  |
| --- | --- | --- | --- |
| **RunUMAP** | dims | umap.method |  |
|  | 1 to 30 | uwot |  |
| **FindNeighbours** | dims | prune.snn | k.param |
|  | 1 to 30 | 1/10 | 40 |
| **FindClusters** | resolution |  |  |
|  | 0,8 |  |  |

| Reclustering of Monocytes and<br>Macrophages cells from WT/CCR2 KO mice<br>data as depicted in Fig 4 |  |  |  |
| --- | --- | --- | --- |
| **RunUMAP** | dims | umap.method |  |
|  | 1 to 30 | uwot |  |
| **FindNeighbours** | dims | prune.snn | k.param |
|  | 1 to 30 | 1/10 | 40 |
| **FindClusters** | resolution |  |  |
|  | 0,7 |  |  |

#### Trajectory inference using Monocle-2

Monocle-2 was used with the raw count matrix of all monocytes and macrophages from WT data (depicted in Figure 2A). Cells were clustered with ‘densityPeak’ method (rho_threshold = 40, delta_threshold = 10). Top 3000 genes by q-value determined using differentialGeneTest() function were used as ‘ordering genes’. Finally, DDRTree dimensionality reduction was performed and cells were ordered using orderCells() function.

#### Inferring cell-cell communication using NicheNet

We used NicheNet (nichenetr R package; version - 0.1.0) to study the interactions between EGCs and infiltrating monocytes after muscularis inflammation. To identify EGC derived ligands potentially inducing differentiation of Ly6c^+^ monocytes into Ccr2^+^ inte Mφs, RiboTag bulk sequencing data from PLP1^CreERT2^ Rpl22^HA^ mice in homeostasis and 3h after the induction of muscularis inflammation were used. Ligands were identified after filtering for genes upregulated 3h post-injury in PLP1^+^ EGCs with respect to homeostasis (adjusted p value < 0.05) and enriched in the HA sample with respect to the IgG control. Genes differentially expressed between Ccr2^+^ int Mφs and Ly6c^+^ monocytes (adjusted p value < 0.05) were considered as the gene set of interest. Genes expressed in at least 10% of the Ly6c+ monocyte cluster excluding genes in the gene set of interest were considered as the background expressed genes in Ly6c^+^ monocytes.

#### Single-cell regulatory network inference and clustering (SCENIC)

SCENIC analysis was run as described by Aibar *et al.* (0.1.5) using the 10-thousand motifs annotation database and 500bp annotation database using pySCENIC (version 0.10.0)^11^ on monocytes and Mφs (2249 cells). The input matrix was the UMI ‘corrected’, log normalized expression matrix. Databases of CisTarget - (mm10 refseq-r80 10kb_up_and_down_tss.mc9nr.feather, mm10 refseq-r80 500bp_up_and_100bp_down_tss.mc9nr.feather), and the transcription factor motif annotation database (motifs-v9-nr.mgi-m0.001-o0.0.tbl) from resources.aertslab.org/cistarget/, and the list of human transcription factors (mm_mgi_tfs.txt) from github.com/aertslab/pySCENIC/tree/master/resources were employed for the analysis. Significance of regulons for each cluster was assessed by a Wilcoxon test on AUC values.

#### Functional analysis of gene sets

Differentially upregulated genes in each cluster compared to the others were used with enrichGO function from the clusterprofiler R package (version 3.14.3)^12^ to perform gene set analysis against the Gene Ontology annotation from org.Mm.eg.db R package (version 3.10.0). Data visualization was prepared using ggplot2 R package.

#### Reference-based annotation using SingleR

SingleR was used for reference-based annotation^13^. Data from Immgen was employed as reference to guide the annotation of CD45^+^ immune cells sorted from the muscularis of WT mice^14^. For the annotation of the mono/Mφ subsets from WT and CCR2^-/-^ mice, the manually curated annotations from the monocyte/Mφ subpopulations of Fig. 2 was used as reference.

#### Gene Set Enrichment Analysis

Gene Set Enrichment Analysis (GSEA) was performed using the GSEA software from the Broad Institute (version 4.0.2). Normalized counts of bulk RNA sequencing from PLP1^+^ cells from the muscularis of PLP1^CreERT2^ Rpl22^HA^ mice at homeostasis and 3h after the induction of muscularis inflammation were used. Only genes that were significantly upregulated in the HA-tagged sample over the IgG control (adjusted p value cut off < 0.05 and log2(fold change) > 0.05) were used for the analysis. Extra data visualization was prepared using ggplot2 R package.

## Supplementary figures

**Supplementary Figure 1:**
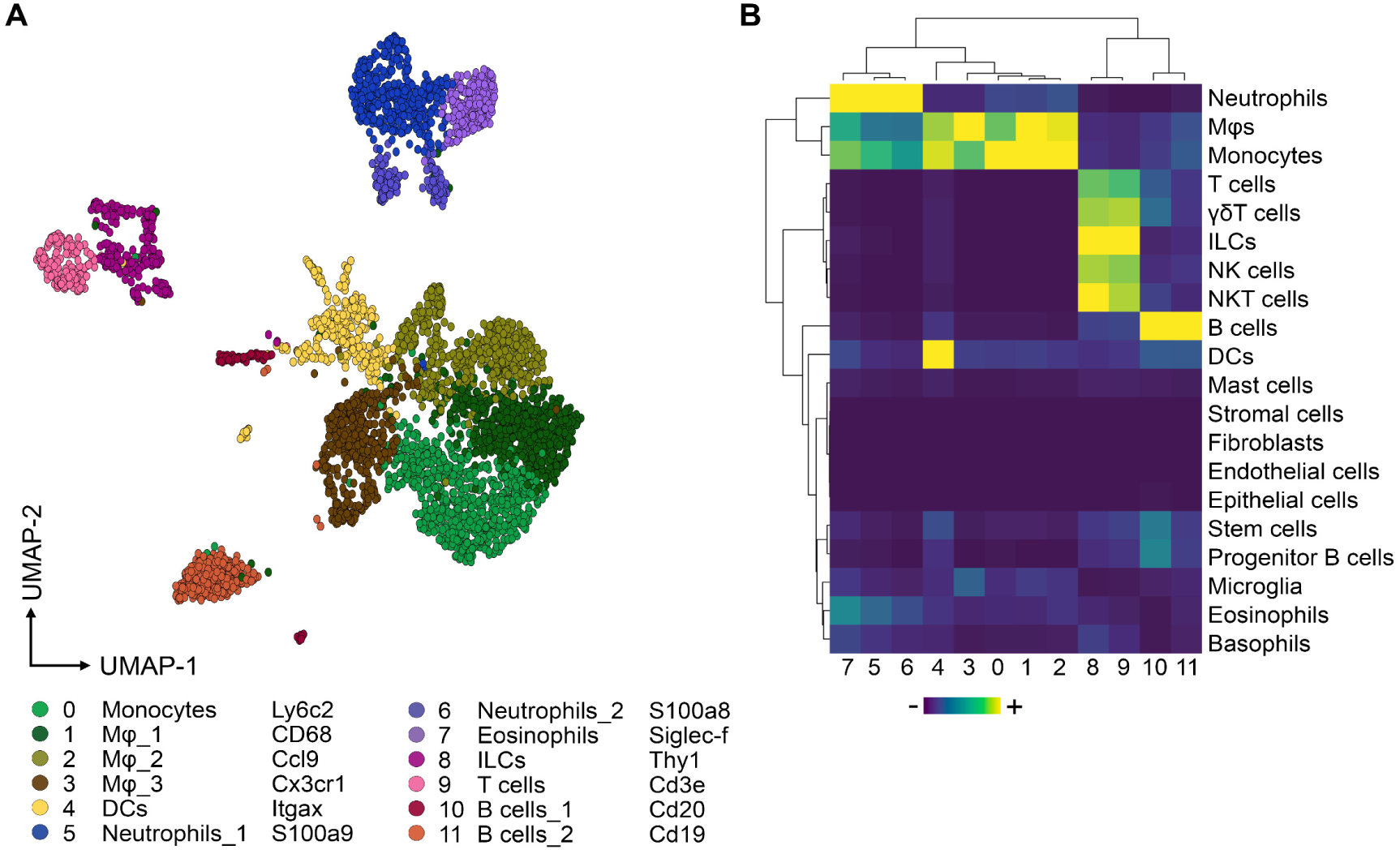
SingleR annotation of CD45^+^ immune cells in the healthy and inflamed muscularis. **A.** UMAP of sorted CD45^+^ immune cells from the healthy muscularis, 24h and 72h after the induction of muscularis inflammation. Reproduction of Figure 1B. **B.** Automated SingleR annotation of clusters from Supplementary Fig. 1A using ImmGen as a reference database.

**Supplementary Figure 2:**
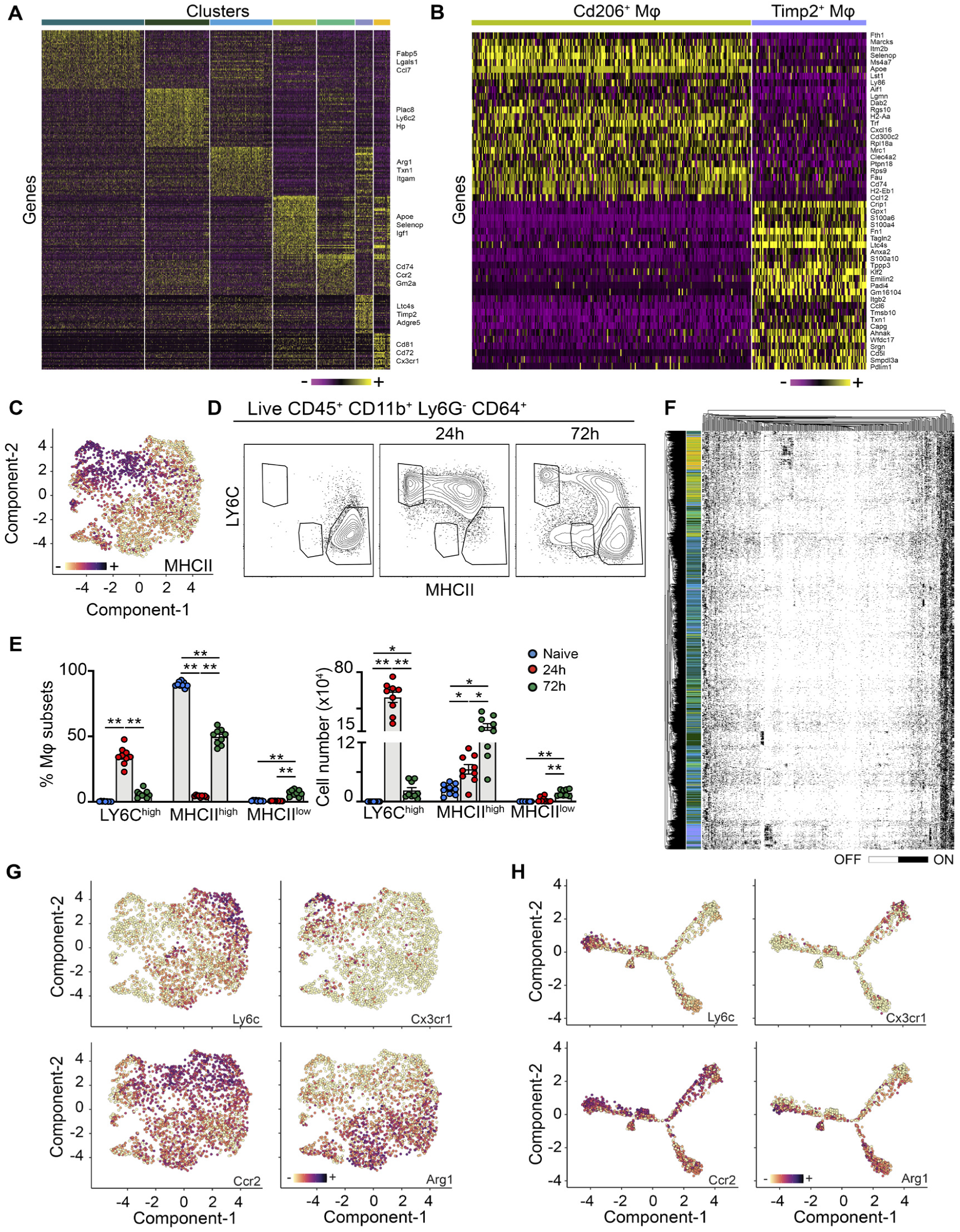
Marker expression in myeloid sub-clustering. **A.** Heatmap of 50 most differentially expressed genes in each cluster from Fig. 2A. **B.** Heatmap of 25 most differentially expressed genes between Cd206^+^ and Timp2^+^ Mφs. **C.** UMAP depicting the expression of the MHCII gene signature. **D-E.** Immune cells isolated from the muscularis at homeostasis, 24h and 72h after muscularis inflammation were analyzed via flow cytometry. Representative contour plots showing Ly6C^hi^ monocytes, and Ly6C^-^MHCII^lo^ Mφs and Ly6C^-^MHCII^hi^ Mφs in the live CD45^+^ CD11b^+^ Ly6G^-^ CD64^+^ population (D). Absolute number of Ly6C^hi^ monocytes, Ly6C^-^MHCII^lo^ Mφs and Ly6C^-^MHCII^hi^ Mφs at different time points after muscularis inflammation. Data are shown as mean ± SEM. One-way ANOVA. *p<0.05; **p<0.01. (E). **F.** Hierarchically clustered heatmap showing ‘binarized’ regulon activity (ON/OFF indicated in Black/White) in each cell. Cells are annotated with the same colors as indicated in Figure 2A. **G.** Single cell gene expression of indicated markers projected onto UMAP. **H.** Single cell gene expression of indicated markers projected onto the Monocle trajectory.

**Supplementary Figure 3:**
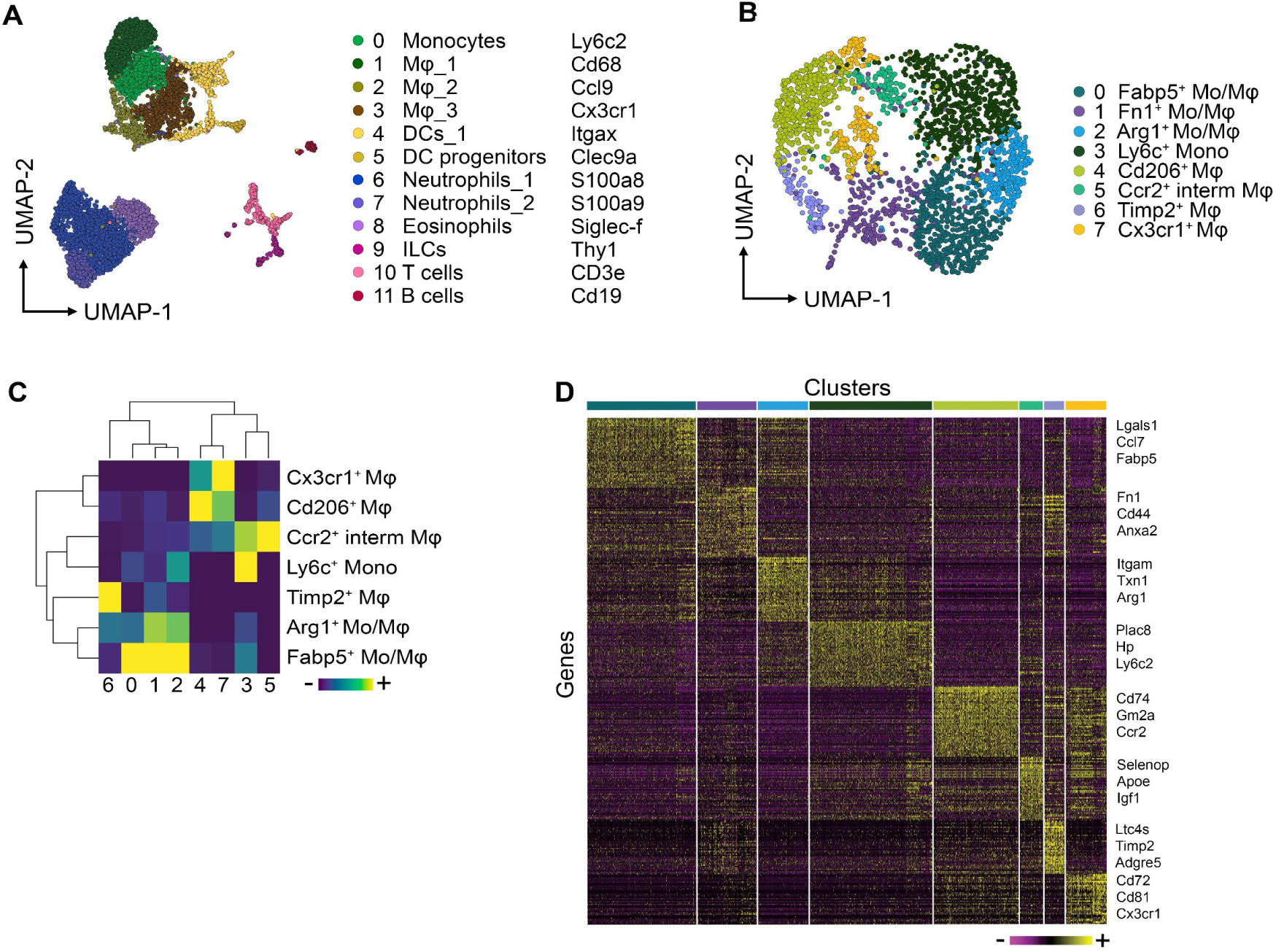
Analysis of myeloid cells in CCR2^-/-^ mice during muscularis inflammation. A-D. scRNAseq on sorted CD45^+^ immune cells from WT and CCR2^-/-^ mice 24h and 72h after the induction of muscularis inflammation. **A.** UMAP of sorted CD45^+^ immune cells. **B.** UMAP of myeloid subclusters. **C.** Automated SingleR annotation of subclusters from Supplementary Fig. 3B based on the gene signature of subclusters from Fig. 2A. **D.** Heatmap of 50 most differentially expressed genes in each subcluster from Supplementary Fig. 3B.

**Supplementary Figure 4:**
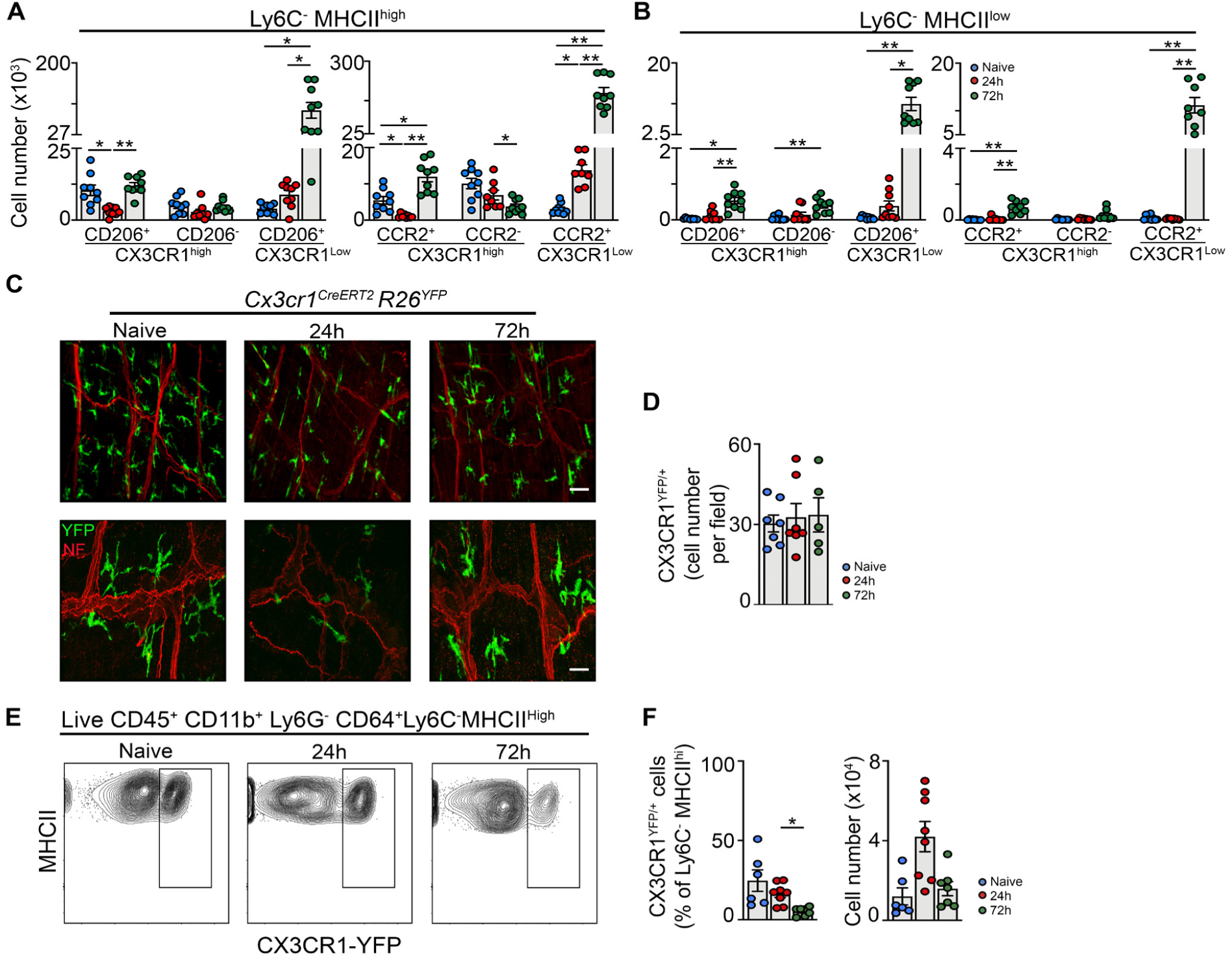
Resident macrophages are not depleted upon muscularis inflammation. **A-B.** Analysis of absolute number of CD206^+^ CX3CR1^hi^, CD206^-^ CX3CR1^hi^ and CD206^+^ CX3CR1^lo^ cells in LY6C^-^ MHCII^hi^ Mφs (A) and LY6C^-^ MHCII^lo^ Mφs (B) and the absolute number of CCR2^+^ CX3CR1^hi^, CCR2^-^ CX3CR1^hi^ and CCR2^+^ CX3CR1^lo^ cells in the LY6C^-^ MHCII^hi^ Mφs (A) and LY6C^-^ MHCII^lo^ Mφs (B) from Fig. 4C,F. **C-F.** Analysis of the number of resident YFP^+^ Mφs during muscularis inflammation (naïve, 24h and 72h) using tamoxifen-injected CX3CR1^CreERT2/+^ Rosa26-LSL-YFP mice. **C.** Representative immunofluorescent images of muscularis whole mount preparations at different time points after muscularis inflammation, stained for YFP (green) and neurofilament (red). Scale bar 40 µm (top; 25X); 15 µm (bottom; 63X). **D.** Number of CX3CR1-YFP^+^ cells per field. Each dot represents the average from an individual mouse and mean ± SEM are shown. Data are combined from 2 independent experiments. **E.** Contour plots showing CX3CR1-YFP expression in LY6C^-^ MHCII^hi^ Mφs at homeostasis and 24h and 72h after muscularis inflammation. **F.** Percentage of CX3CR1^YFP/+^ cells in LY6C^-^ MHCII^hi^ Mφs (left) and absolute number of cells (right) at different time points after muscularis inflammation. A, B, D, F; one-way ANOVA. *p<0.05; **p<0.01, ns= not significant.

**Supplementary Figure 5:**
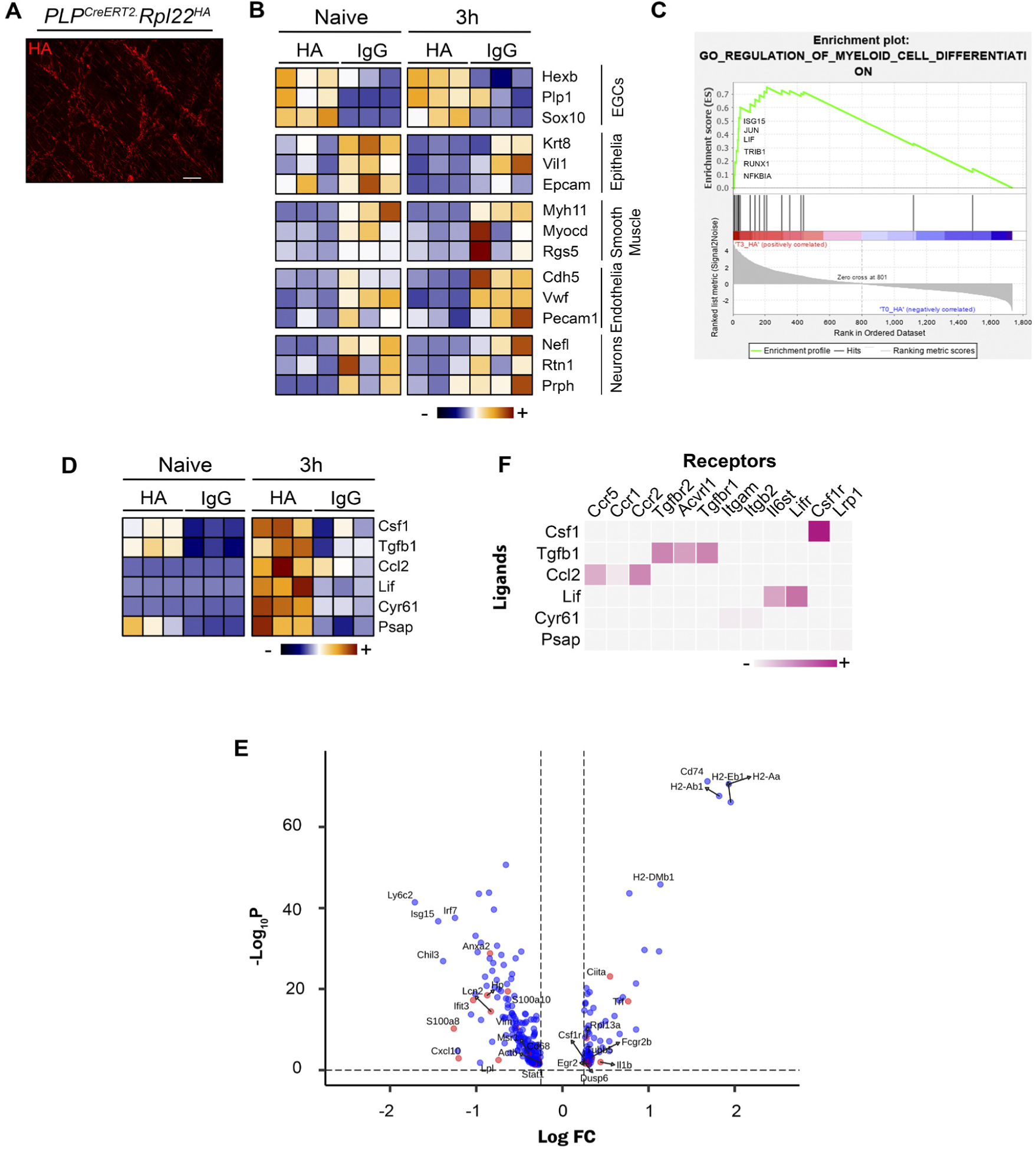
Characterization of EGCs at homeostasis and during muscularis inflammation. **A.** Immunofluorescent image of muscularis whole mount preparation of a tamoxifen-injected PLP^CreERT2/+^ Rpl22^HA^ mouse stained for HA tag (red). Scale bar 30 µm. **B.** Heat map of gene expression from IgG control or immunoprecipitated muscularis from naive mice or 3h after the induction of muscularis inflammation showing genes characteristics for different cell types. **C.** Visualization of GSEA analysis for regulation of myeloid cell differentiation. **D.** Heatmap of all positively correlated upstream ligands from NicheNet analysis. **E.** Volcano plot of differentially expressed genes between Ccr2^+^ int Mφs and Ly6c^+^ monocytes highlighting target genes of top ligands from NicheNet in Ly6c^+^ monocytes and Ccr2^+^ int Mφs. **F.** Heatmap showing prior interaction potential between the ligands and corresponding receptors.

**Supplementary Figure 6:**
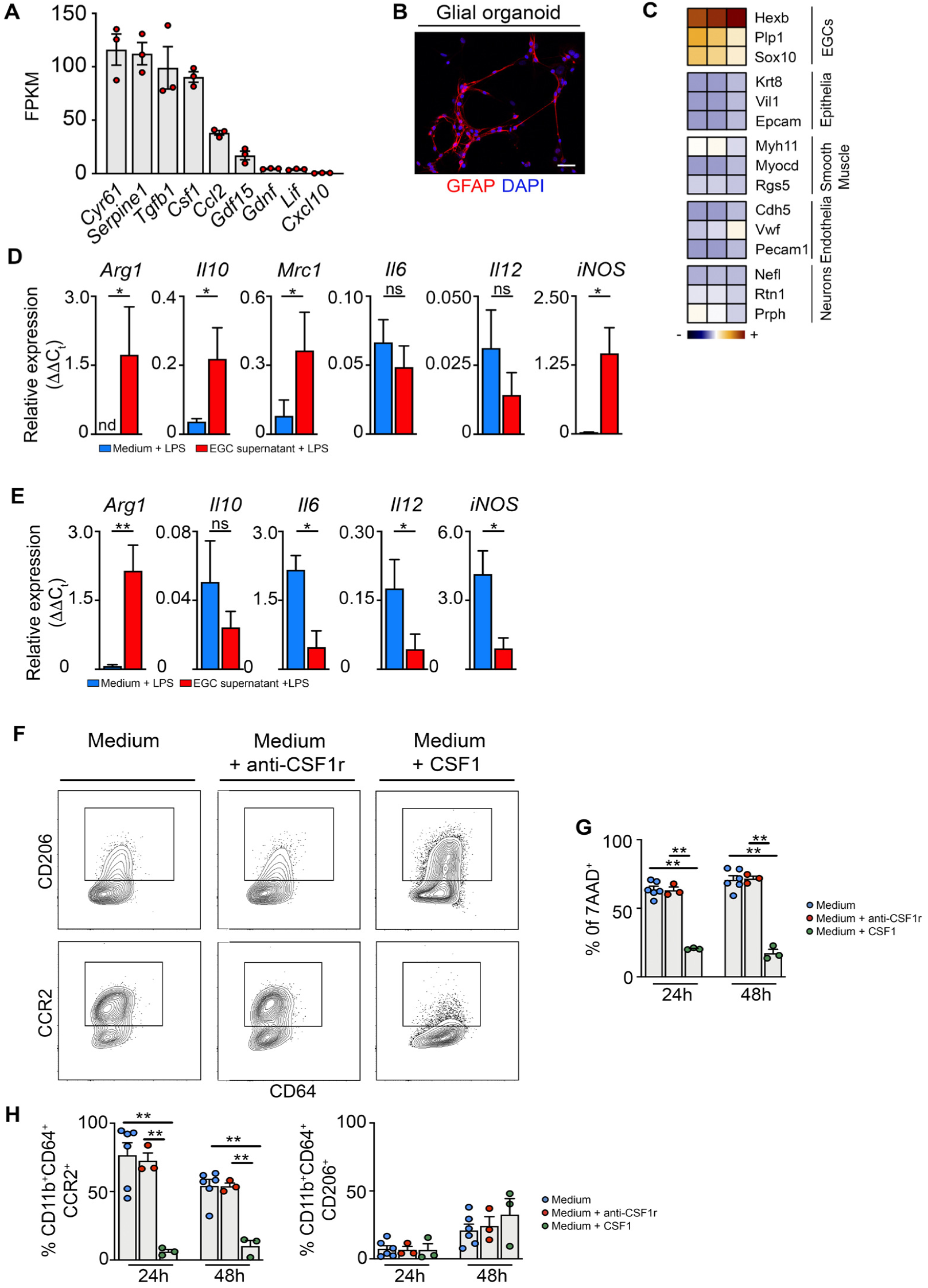
Supernatant of neurosphere-derived EGCs blunts inflammatory response in monocytes upon stimulation with LPS. **A.** FPKM values of RNAseq performed on unstimulated neurosphere-derived EGCs. **B.** Neurosphere-derived EGCs stained with GFAP (red) and DAPI (blue). Scale bar 40 µm. **C.** BM monocytes were cultured for 18h with/without EGC supernatant followed by LPS stimulation (100 ng/mL) for 6h. Relative mRNA levels for pro- and anti-inflammatory cytokines normalized to the housekeeping gene *rpl32* are shown. **D.** Relative mRNA levels for pro- and anti-inflammatory cytokines normalized to the housekeeping gene *rpl32* in Ly6C^hi^ monocytes from the muscularis 24 post-injury and cultured 18h with or without EGC supernatant and with LPS (100 ng/mL) for 6h. **E-G.** BM monocytes were cultured for 24-48h with medium alone, supplemented with anti-CSF1r (5 µg/mL) or CSF-1 (50 ng/mL). **E.** Contour plots of the expression of CD206 (top) and CCR2 (bottom) at 48h of culture. **F.** Quantification of 7-AAD^+^ cells in BM monocytes cultured for 24h and 48h. **G.** Percentage of CCR2^+^ and CD206^+^ cells in the live CD45^+^ CD11b^+^ Ly6G^-^ CD64^+^ population. C-D. t-test. F-G; one-way ANOVA. *p<0.05; **p<0.01.

